# LeafCutter: annotation-free quantification of RNA splicing

**DOI:** 10.1101/044107

**Authors:** Yang I Li, David A Knowles, Jack Humphrey, Alvaro N. Barbeira, Scott P. Dickinson, Hae Kyung Im, Jonathan K Pritchard

**Author notes:** These authors contributed equally to this work (Yang I Li, David A Knowles). Correspondence should be addressed to Y.I.L., D.A.K., or J.K.P.

## Abstract

The excision of introns from pre-mRNA is an essential step in mRNA processing. We developed LeafCutter to study sample and population variation in intron splicing. LeafCutter identifies variable intron splicing events from short-read RNA-seq data and finds alternative splicing events of high complexity. Our approach obviates the need for transcript annotations and circumvents the challenges in estimating relative isoform or exon usage in complex splicing events. LeafCutter can be used both for detecting differential splicing between sample groups, and for mapping splicing quantitative trait loci (sQTLs). Compared to contemporary methods, we find 1.4–2.1 times more sQTLs, many of which help us ascribe molecular effects to disease-associated variants. Strikingly, transcriptome-wide associations between LeafCutter intron quantifications and 40 complex traits increased the number of associated disease genes at 5% FDR by an average of 2.1-fold as compared to using gene expression levels alone. LeafCutter is fast, scalable, easy to use, and available at https://github.com/davidaknowles/leafcutter.

## Background

The alternative removal of introns during mRNA maturation is essential for major biological processes in eukaryotes including cellular differentiation, response to environmental stress, and proper gene regulation ^1,2,3,4^. Nevertheless, our ability to draw novel insights into the regulation and function of splicing is hindered by the challenge of estimating transcript abundances from short-read RNA-seq data.

Most popular approaches for studying alternative splicing from RNA-seq estimate isoform ratios ^5,6,7,8^ or exon inclusion levels ^9,10^. Quantification of isoforms or exons is intuitive because RNA-seq reads generally represent mature mRNA molecules from which introns have already been removed. However, estimation of isoform abundance from conventional short-read data is statistically challenging, as each read samples only a small part of the transcript, and alternative transcripts often have substantial overlap ^11^. Similarly, when estimating exon expression levels from RNA-seq data, read depths are often overdispersed due to technical effects, and there may be ambiguity about which version of an exon is supported by a read if there are alternative 5’ or 3’ splice sites.

Further, both isoform-and exon-quantification approaches generally rely on transcript models, or predefined splicing events, both of which may be inaccurate or incomplete ^12^. Predefined transcript models are particularly limiting when comparing splicing profiles of healthy versus disease samples, as aberrant transcripts may be disease-specific; or when studying genetic variants that generate splicing events in a subset of individuals only ^13^. Even when transcript models are complete, it is difficult to estimate isoform or exon usage of complex alternative splicing events ^12^.

An alternative perspective is to focus on what is removed in each splicing event. Excised introns may be inferred directly from reads that span exon-exon junctions. Thus, there is generally little ambiguity about the precise intron that is cut out, and quantification of usage ratios is very accurate ^12^. A recent method, MAJIQ ^12^, also proposed to estimate local splicing variation using split-reads and identified complex splicing events, however it does not scale well above 30 samples and has not been adapted to map splicing QTLs (sQTLs). At present, there are several software programs for sQTL mapping: GLiMMPS ^14^, sQTLseekeR ^15^ and Altrans ^16^. However, all three rely on existing isoform annotations and both GLiMMPS and sQTLseeker reported modest numbers of sQTLs in their analyses.

Here we describe LeafCutter, a suite of novel methods that allow identification and quantification of novel and existing alternative splicing events by focusing on intron excisions. We show LeafCutter’s utility by applying it to three important problems in genomics: (1) identification of differential splicing across conditions, (2) identification of sQTLs in multiple tissues or cell types, and (3) ascribing molecular effects to disease-associated GWAS loci. Using an early version of LeafCutter, we found that alternative splicing is an important mechanism through which genetic variants contribute to disease risk ^17^. We now show that LeafCutter dramatically increases the number of detectable associations between genetic variation and pre-mRNA splicing, thus enhancing our understanding of disease-associated loci.

## Results

### Overview of LeafCutter

LeafCutter uses short-read RNA-seq data to detect intron excision events at base-pair precision by analyzing split-mapped reads. LeafCutter focuses on alternative splicing events including skipped exons, 5’ and 3’ alternative splice site usage and additional complex events that can be summarized by differences in intron excision ^12^ (Supplementary Figure 1). LeafCutter’s intron-centric view of splicing is motivated by the observation that mRNA splicing predominantly occurs through the step-wise removal of introns from nascent pre-mRNA ^19^. (Unlike isoform quantification methods such as Cufflinks2 ^5^, alternative transcription start sites, and alternative polyadenylation are not directly measured by LeafCutter as they are not generally captured by intron excision events.) The major advantage of this representation is that LeafCutter does not require read assembly or inference on which isoform is supported by ambiguous reads, both of which are computationally and statistically difficult problems. An implication of this is that we were able to improve speed and memory requirements by an order of magnitude or more as compared to similar methods such as MAJIQ^12^.

To identify alternatively-excised introns, LeafCutter pools all mapped reads from a study and finds overlapping introns demarcated by split reads. LeafCutter then constructs a graph that connects all overlapping introns that share a donor or an acceptor splice site (Figure 1a). The connected components of this graph form clusters, which represent alternative intron excision events. Finally, LeafCutter iteratively applies a filtering step to remove rarely used introns, which are defined based on the proportion of reads supporting an intron compared to other introns in the same cluster, and re-clusters leftover introns (Methods, Supplementary Note 1). In practice, we found that this filtering is important to avoid arbitrarily large clusters when read depth increases to a level at which noisy splicing events are supported by multiple reads.

**Figure 1:**
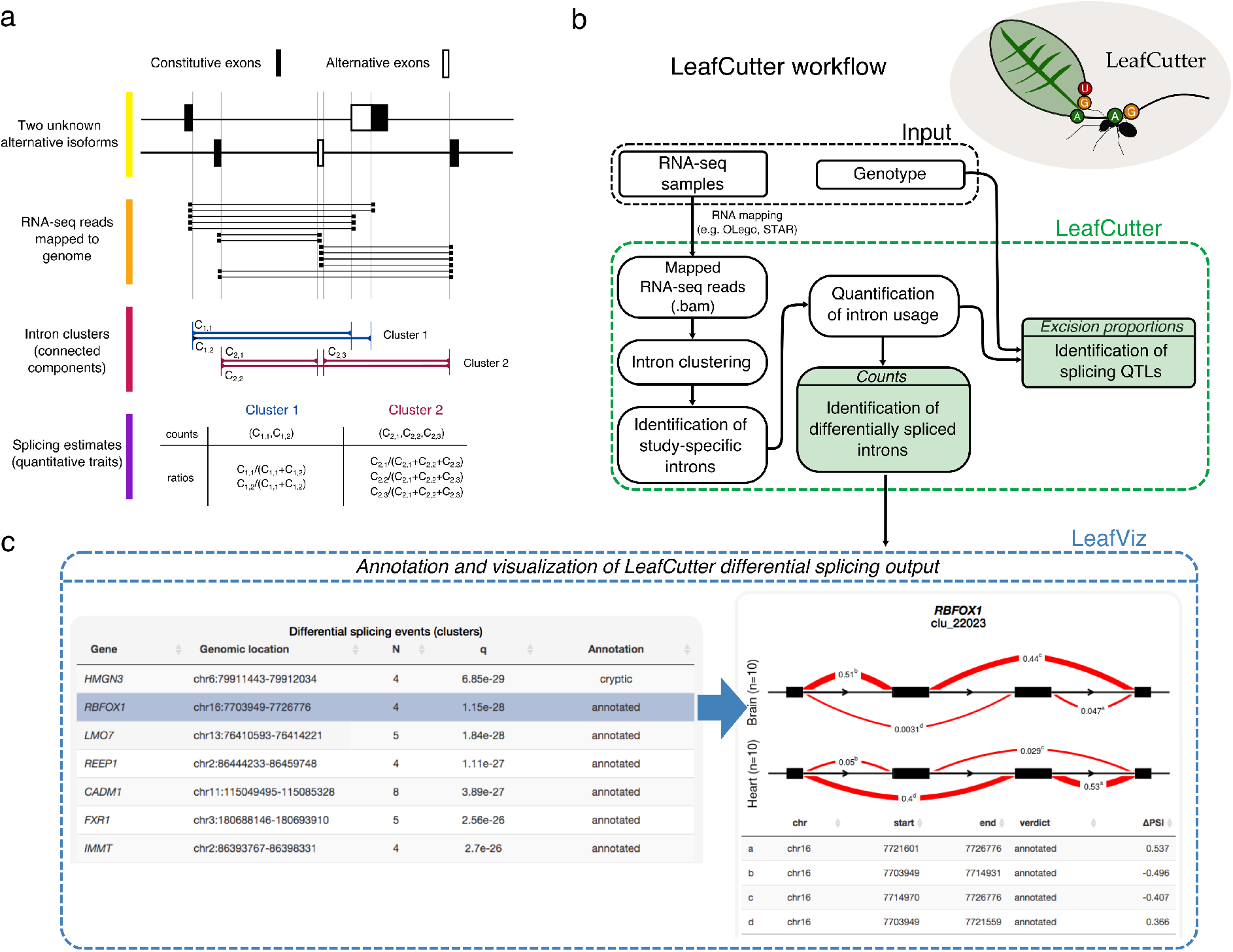
Overview of LeafCutter. (a) LeafCutter uses split reads to uncover alternative choices of intron excision by finding introns that share splice sites. In this example, LeafCutter identifies two clusters of variably excised introns. (b) LeafCutter workflow. First, short reads are mapped to the genome. When SNP data are available, WASP ^18^ should be used to filter allele-specific reads that map with a bias. Next, LeafCutter extracts junction reads from .bam files, identifies alternatively excised intron clusters, and summarizes intron usage as counts or proportions. Lastly, LeafCutter identifies intron clusters with differentially excised introns between two user-defined groups using a Dirichlet-multinomial model or maps genetic variants associated with intron excision levels using a linear model. (c) Visualization of differential splicing between 10 GTEx heart and brain samples using LeafViz. LeafViz is an interactive browser-based application that allows users to visualize results from LeafCutter differential splicing analyses. In this example, we observed that *Rbfox1* shows differential usage of a mutually exclusive exon in heart compared to brain. For all examples, visit https://leafcutter.shinyapps.io/leafviz/.

### *De novo* identification of functional RNA splicing in mammalian organs

We tested LeafCutter’s novel intron detection method by analyzing mapped RNA-seq ^20^ short read data from 2,192 samples (Supplementary Note 3) across 14 tissues from the GTEx consortium ^21^. We then searched introns predicted to be alternatively excised by LeafCutter, but that were missing in three commonly-used annotation databases (GENCODE v19, Ensembl, and UCSC). For this analysis, we ensured that the identified introns were indeed alternatively excised by only considering introns that were excised at least 20% of the time as compared to other overlapping introns, in at least one fourth of the samples, analyzing each tissue separately. We found that between 10.8% and 19.3% (Pancreas and Spleen, respectively) of alternatively spliced introns are unannotated – excluding testis which is the major outlier, in which 48.5% of alternatively spliced introns are novel (Figure 2a). The latter observation is compatible with the “out-of-testis” hypothesis, which proposes that transcription is more permissive in testis and allows novel genes or isoforms to be selected for if beneficial ^22,23^. Thus 31.5% of the alternatively excised introns we detected are unannotated (Supplementary Note 4), consistent with a recent study that identified a similar proportion of novel splicing events in 12 mouse tissues ^12^. To further confirm that these findings were not merely mapping or GTEx-specific artefacts, we searched for junction reads in 21,504 human RNA-seq samples from the Sequence Read Archive (SRA) obtained from `Intropolis` ^24^. We found that most (86%, Figure 2c and Supplementary Figure S13) novel junctions identified in our study were also present in at least one RNA-seq sample from the corresponding tissue as identified in `Intropolis`. Furthermore, we found that, as expected, unannotated junctions tend to be tissue-specific, and often involve complex splicing patterns (Supplementary Figure S14 and Supplementary Note 4).

**Figure 2:**
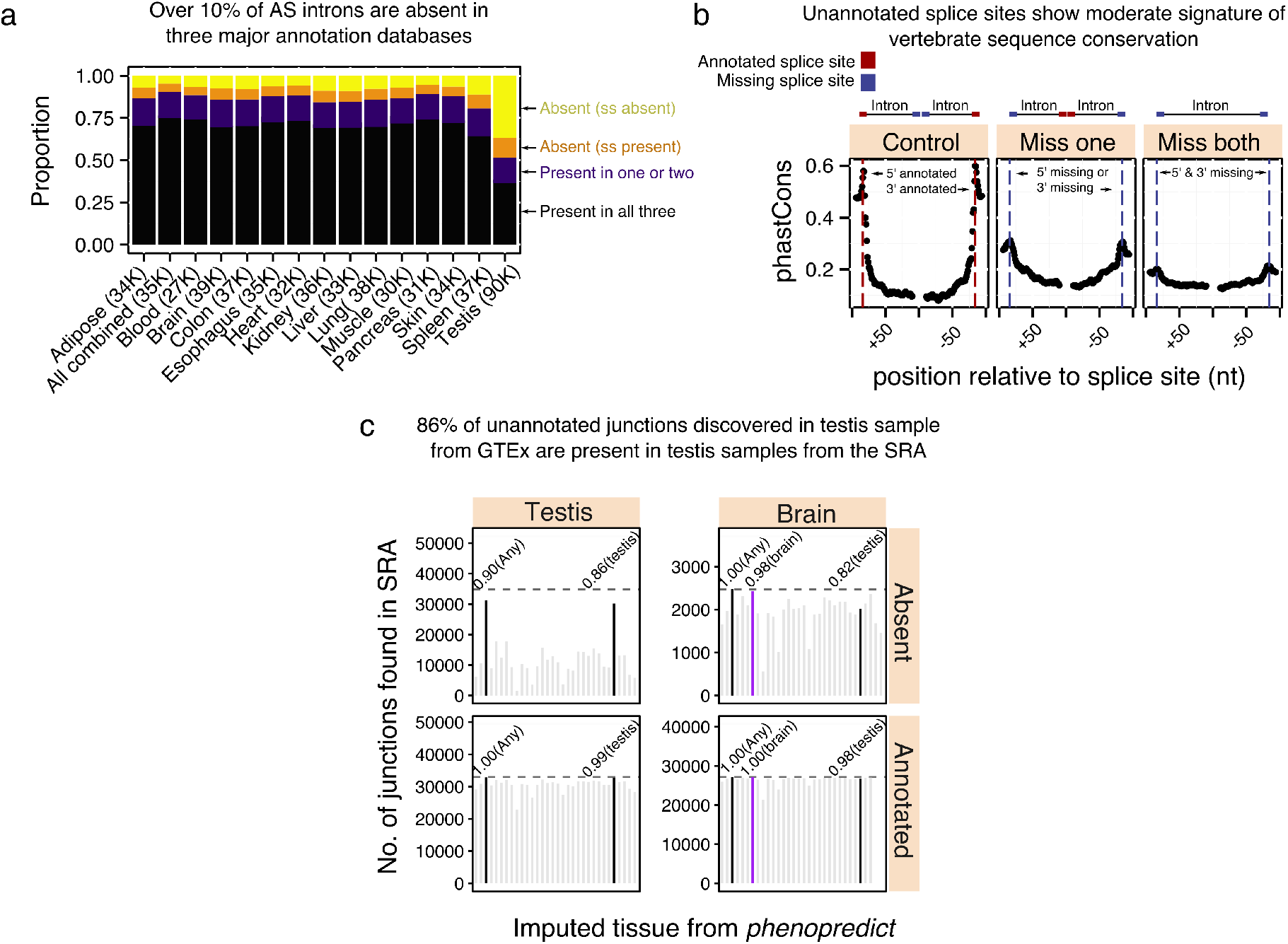
(a) Using LeafCutter to discover novel introns, we find that for any given tissue, over 10% of alternatively excised introns are unannotated. Remarkably, 48.5% of testis alternatively excised introns are unannotated. Different colors denote the proportion of introns when one or more splice sites are unannotated “(ss absent)”, both splice sites are annotated but the intron is not part of any transcript “(ss present)”, or when the intron is annotated in some but not all databases. (b) The unannotated splice sites of novel introns show moderate signature of sequence conservation as determined by vertebrate phastCons scores. Miss one: conservation of the unannotated splice site of an intron for which the cognate splice site is annotated. Miss both: conservation of splice sites of introns with both splice sites unannotated. (c) Barplots showing the numbers of unannotated and annotated junctions discovered using LeafCutter that are also found in samples from the short read archive (SRA) using `Intropolis` ^24^. `Phenopredict` ^25^ was used to predict the tissue type corresponding to the SRA samples analyzed in `Intropolis`.

We next asked whether these novel introns show evidence of functionality as determined by sequence conservation. When we averaged phastCons scores over unannotated splice sites of introns that were absent in annotation databases, we found a moderate, but significant, signature of sequence conservation (Figure 2b). In particular, we found that a significant number (4,616 or 15–25%) of novel splice sites are conserved across vertebrates (ave. phastCons ≥ 0.6, Supplementary Figure S11), indicating that the alternative excision of thousands of introns may be functional (Supplementary Note 4).

### Fast and robust identification of differential splicing across sample groups

LeafCutter uses counts from the clustering step (Figure 1b) to identify introns with differential splicing between user-defined groups. Read counts in an intron cluster are jointly modeled using a Dirichlet-multinomial generalized linear model (GLM), which we found offers superior sensitivity relative to a beta-binomial GLM that tests each intron independently (Supplementary Figure S15). The implicit normalization of the multinomial likelihood avoids the estimation of library size parameters required by methods such as DEXSEQ ^10^.

We compared LeafCutter against other methods for differential splicing detection including Cufflinks2 ^5^, MAJIQ ^12^, and rMATS ^26^. We note that comparisons between algorithms have the complication that they there is typically no one-to-one mapping between the splicing events quantified by different methods. We discuss this issue and our solution in Supplementary Note 2. For comparison, we applied each method to identify splicing differences between 3, 5, 10, and 15 Yoruba (YRI) versus European (CEU) LCL RNA-seq samples. In terms of runtime, we observed a large difference in scalability (Figure 3a). In our hands, only LeafCutter completed all comparisons within an hour, while Cufflinks2, rMATS, and MAJIQ took as long as 7.8, 55.7, and 66.2 hours to complete the largest comparison, respectively. In terms of memory usage, we also found that LeafCutter greatly outperforms the other software, using less than 400Mb of RAM for all comparisons, while MAJIQ required over 50Gb of RAM to perform the larger comparisons (Supplementary Figure S9). Although this range of sample sizes is representative for most biological studies, identifying differential splicing across groups in large studies such as GTEx would be impractically slow using rMATS or MAJIQ.

**Figure 3:**
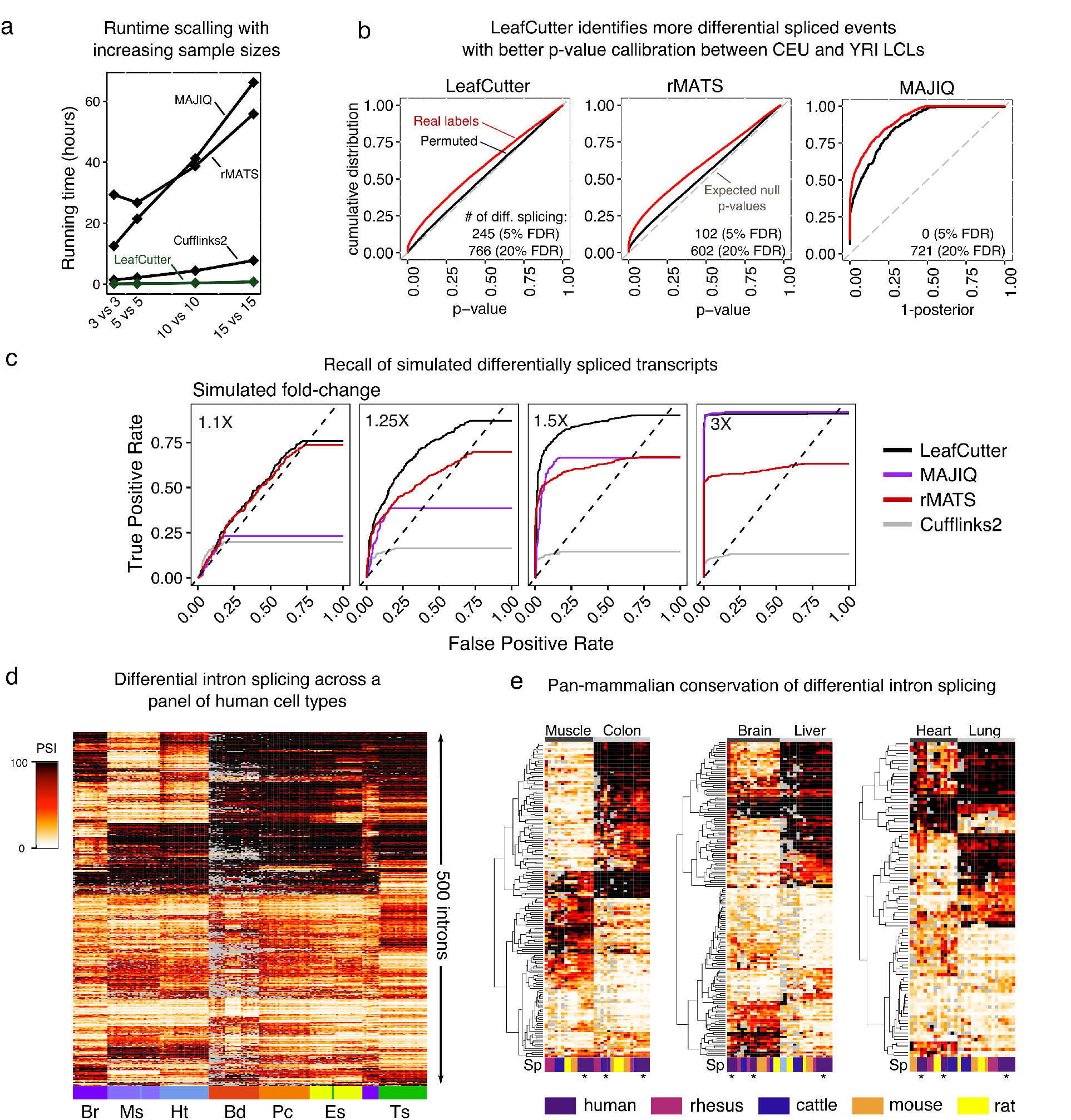
Comparison of LeafCutter against other methods for detecting differential splicing. (a) Running time of various differential splicing methods applied to comparisons between 3, 5, 10, and 15 YRI vs CEU LCLs RNA-seq samples. (b) Cumulative distributions of differential splicing test p-values (1-posterior for MAJIQ) for the 15 YRI versus 15 CEU LCLs comparison (red). The distribution of test p-values for a 15 versus 15 samples comparison with permuted labels is also shown (black). LeafCutter detects more differential splicing compared to rMATS, MAJIQ, and Cufflinks2. Cufflinks2 distribution omitted as it detected 0 significantly differentially spliced genes as described in Supplementary Note 2.3 (Supplementary Figure S3). (c) Receiver operating characteristic (ROC) curves of LeafCutter, Cufflinks2, rMATS and MAJIQ when evaluating differential splicing of genes with transcripts simulated to have varying levels of differential expression. ROC curves that do not reach 1.00 True Positive Rates reflect the proportion of genes simulated to be differentially spliced that were not tested (Supplementary Note 2.4). (d) LeafCutter identifies tissue-regulated intron splicing events from GTEx organ samples. Heatmap of the intron excision ratios of the top 500 introns that were found to be differentially spliced between at least one tissue pair. Tissues include brain (Br), muscle (Ms), heart (Ht), blood (Bd), pancreas (Pc), esophagus (Eg), and testis (Ts). (e) Tissue-dependent intron excision is conserved across mammals. Heatmap showing intron exclusion ratios of introns differentially spliced between pairs of tissues (Muscle vs Colon, Brain vs Liver, and heart vs Lung). Heatmap shows 100 random introns (97 for the heart vs lung comparison) that were predicted to be differentially excised in human with p-value < 10^-10^ (LR-test) and that had no more than 5 samples where the excision rate could not be determined due to low count numbers. Heatmap of all introns that pass our criteria can be found in Supplementary Figure S18.

To compare their ability to detect differential splicing, we reasoned that the *p*-values or posterior probabilities of the tests computed by each method are not directly comparable. We therefore computed an empirical FDR from the *p*-values of real comparisons between biologically distinct sample groups (i.e. YRI versus CEU here) and the *p*-values of permuted comparisons between samples with permuted labels (i.e. both groups contain YRI and CEU samples). If the *p*-values are well-calibrated, the *p*-value distribution of the permuted comparisons are expected to be uniform. Indeed, we observed that the distributions of Leaf-Cutter and rMATS *p*-values for the comparisons were close to the theoretical null distribution (Figure 3b and Supplementary Figure S3). However, we observed that the Cufflinks2 p-values were overly conservative (Supplementary Note 2.3) and additionally the posterior probabilities *P* reported by MAJIQ for the permuted comparisons did not track the expected false discovery rate (FDR) of 1 – *P* (Supplementary Note 2.2). Altogether, we found that LeafCutter *p*-values showed better callibration compared to other methods, and that LeafCutter detected more differentially spliced events at all reasonable FDRs (≤ 0.2). Importantly, not only did LeafCutter detect more differentially spliced events at fixed FDRs, but it also achieved lower false negative rates when we evaluated the four methods on artificial data in which we simulated various levels of fold-changes in isoform levels (Figure 3c, Supplementary Note 2.4, Supplementary Figure S6). These comparisons show that LeafCutter is a robust and highly scalable method for differential splicing analysis.

To evaluate LeafCutter’s suitability to detect differential splicing in a biological setting, we searched for intron clusters that show differential splicing between tissue pairs collected by the GTEx consortium, using all tissues to identify intron clusters. Combining all pairwise comparisons, we found 5,070 tissue-regulated splicing clusters at 10% FDR and with an estimated absolute effect size greater than 1.5 (Methods). As expected, GTEx samples mostly grouped by organ/tissue when hierarchically clustered according to the excision ratios of the five hundred most differentially spliced introns among all tissue pairs (Figure 3d, Supplementary Note 5).

To evaluate LeafCutter’s applicability to studies with smaller sample sizes, we used LeafCutter on a small subset of all available GTEx samples and then evaluated the amount of replication when using a larger subset. When using 220 samples (110 brain versus 110 muscle samples), we identified 1,906 differentially spliced clusters with estimated effect sizes greater than 1.5 at 10% FDR, compared to 885 when using only 8 samples (4 brain versus 4 muscle samples). Importantly, the strengths of associations (-log_10_ *p*-values) were highly correlated between our two analyses (Pearson R^2^ = 0.72, Supplementary Note 6, Supplementary Figure S17), and 98% of alternatively spliced clustered identified at 10% FDR in the analysis using 8 total samples were replicated in the analysis using 220 samples, also at 10% FDR. These observations indicate that LeafCutter can be used to detect differentially spliced introns even when the number of biological replicates is small.

We then investigated whether the differentially spliced clusters identified using LeafCutter are likely to be functional by assessing the pan-mammalian conservation of their splicing patterns across multiple organs. Two previous studies analyzed the evolution of alternative splicing in mammals and, when they clustered samples using gene expression levels, saw clustering by organ as expected, however when they clustered samples using exon-skipping levels, they instead saw a clustering by species ^27,28^. These observations indicate that a large number of alternative skipping events may lack function or undergo rapid turnover.

We initially attempted clustering using all splicing events and confirmed the previous findings ^27,28^ that the samples mostly clustered by species (Supplementary Figure S16). We then focused on a subset of introns that LeafCutter identified as differentially excised across tissue pairs in human and found that this subset shows splicing patterns that are broadly conserved across mammalian organs (Figure 3e). To do this, we hierarchically clustered samples from eight organs in human and four mammals ^27^ according to the orthologous intron excision proportions of differentially excised introns (*p*-value < 10^-10^ and *β* > 1.5) from our pairwise analyses of human GTEx samples (Supplementary Methods 5). Unlike in the previous analyses, this revealed a striking clustering of the samples by organ, implying that hundreds of tissue-biased intron excisions events are conserved across mammals and likely have organ-specific functional roles ^29^. Thus, while the majority of alternative splicing events likely undergo rapid turnover, events that show organ-specificity are much more often conserved across mammals and, therefore, are more likely to be functionally important.

### Mapping splicing QTLs using LeafCutter

Next, to evaluate LeafCutter’s ability to map splicing QTLs, we applied LeafCutter to 372 EU lymphoblastoid cell line (LCL) RNA-seq samples from GEUVADIS, and identified 42,716 clusters of alternatively excised introns. We used the proportion of reads supporting each alternatively excised intron identified by LeafCutter and a linear model ^30^ to map sQTLs (Supplementary Note 7). We found 5,774 sQTLs at 5% FDR (compared to 620 trQTLs in the original study at 5% FDR, i.e., one ninth times as many) and 4,543 at 1% FDR. To perform a controlled comparison, we also processed 85 YRI GEUVADIS LCLs RNA-seq samples and quantified RNA splicing events using LeafCutter, Altrans ^16^, and Cufflinks2 ^5^. We then uniformly standardized and normalized the estimates and used them as input to fastQTL ^30^ to identify sQTLs (Supplementary Note 3.3 and 7.2). At similar false discovery rates, LeafCutter identifies 1.36X–1.46X and 1.83X–2.06X more sQTLs than Cufflinks2 and Altrans, respectively (Table 1). The rate of sQTL discoveries shared between methods is generally high (Storey’s π_1_ ranging from 0.53 to 0.72 for sQTLs identified at 10% FDR, Supplementary Note 7.3, Supplementary Figure S19), with LeafCutter sQTLs showing higher estimates of sharing (π1 = 0.70 and 0.72 with Cufflinks2 and Altrans, respectively) than Cufflinks2 sQTLs (0.52 with Altrans) or Altrans sQTLs (0.66 with Cufflinks2).

**Table 1:** Summary of sQTLs identified in GEUVADIS samples using LeafCutter, Altrans ^16^, and Cufflinks2^5^. The numbers of transcript ratio QTLs (trQTLs) identified in the orginal GEUVADIS study ^31^ are also listed in the sQTL columns. N/A: Not available.

| Method | YRI sQTLs (0.01 FDR) | YRI sQTLs (0.05 FDR) | CEU sQTLs (0.05 FDR) |
| --- | --- | --- | --- |
| LeafCutter | 1,294 | 1,982 | 5,775 |
| Altrans | 624 | 1,083 | N/A |
| Cufflinks2 | 888 | 1,459 | N/A |
| GEUVADIS study | N/A | 83 | 620 |

To further ensure that our sQTLs are not simply false positives, we verified that LeafCutter finds stronger associations between intronic splicing levels and SNPs previously identified as exon eQTLs and transcript ratio QTLs in GEUVADIS ^31^ when compared to genome-wide SNPs (Figure 4a). Importantly, 399 (81.3%) of the 491 top trQTLs tested are significantly associated to intron splicing variation, as identified by LeafCutter (compared to 4.7% when our samples are permuted, Supplementary Note 7). Furthermore, we confirmed that the sQTLs we identified are located near splice sites, are close to the introns they affect (Figure 4b), and are enriched in expected functional annotations such as “splice regions” and DNaseI hypersensitivity regions (Supplementary Figure S22).

**Figure 4:**
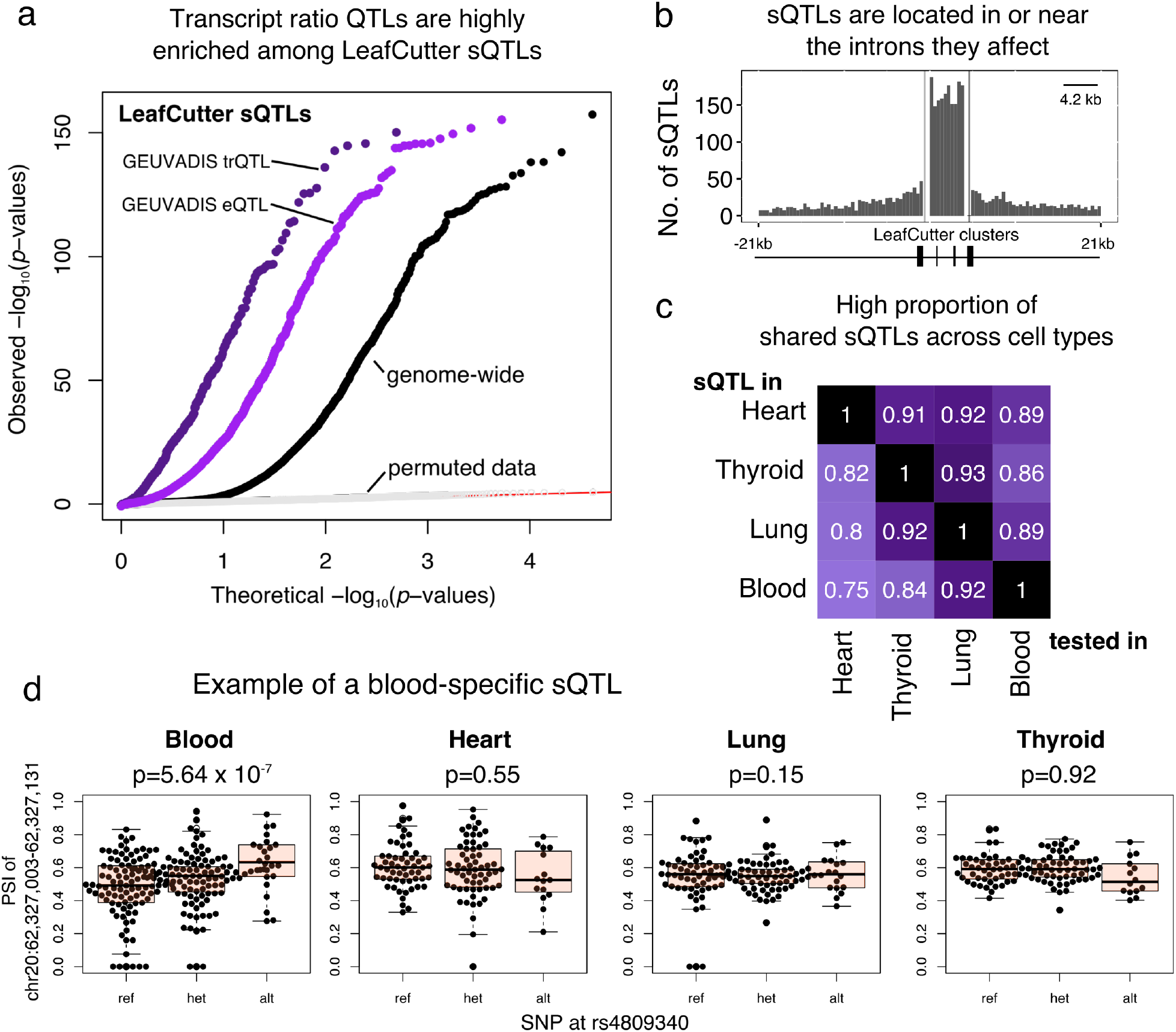
(a) QQ-plot showing genome-wide sQTL signal in LCLs (black), sQTL signal conditioned on exon eQTLs (purple) and conditioned on transcript ratio QTLs (dark purple) from ^31^. Signal from permuted data in light grey shows that the test is well-calibrated. (b) Positional distribution of sQTLs across LeafCutter-defined intron clusters. 1,421 of 4,543 sQTLs lie outside the boundaries (Supplementary Figure S22 for all sQTLs). (c) High proportion of shared sQTLs across four tissues from ^21^. (d) Example of a SNP associated to the excision level of an intron in blood but not in other tissues.

We used LeafCutter to identify sQTLs in four tissues from the GTEx consortium. Overall, we found 442, 1,058, 1,047, and 692 sQTLs at 1% FDR in heart, lung, thyroid gland, and whole blood, respectively (Supplementary Note 7). Using these, we estimated that 75–93% of sQTLs replicate across tissue pairs (Figure 4c, Supplementary Figure S24, Supplementary Note 6). This agrees with a high proportion of sharing of sQTLs across tissues ^32^; and contrasts with much lower pairwise sharing reported for these data previously (9-48%) ^21^. The high level of replication is likely owing to LeafCutter’s increased power in detecting genetic associations with specific splicing events. Nevertheless, this leaves 7-25% of sQTLs that show tissue-specificity in our analysis. As expected we found that a large proportion of tissue-specific sQTLs arose from trivial cases where the intron is only alternatively excised, and therefore variable, in one tissue (Supplementary Figure S25). However, we also found cases in which the introns were alternatively excised in all tissues, yet show tissue-specific association with genotype (Figure 4d).

### LeafCutter sQTLs link disease-associated variants to mechanism

Finally, we asked whether sQTLs identified using LeafCutter could be used to ascribe molecular effects to disease-associated variants as determined by genome-wide association studies. For example, eQTLs are enriched for disease-associated variants, and disease-associated variants that are eQTLs likely function by modulating gene expression ^31,21^. We recently showed that sQTLs identified in LCLs are also enriched among autoimmune-disease-associated variants ^17^. LeafCutter sQTLs can therefore help us characterize the functional effects of variants associated with complex diseases. Indeed, when we looked at the association signals of the top LeafCutter sQTLs and eQTLs from GEUVADIS to multiple sclerosis and rheumatoid arthritis (Supplementary Note 8), we found that both QTL types were enriched for stronger associations (Figure 5a) compared to genome-wide variants. Consistent with recent findings ^17^, SNPs associated with multiple sclerosis are more highly enriched among sQTLs than eQTLs, while both eQTLs and sQTLs are similarly enriched among SNPs associated with rheumatoid arthritis (Figure 5a).

**Figure 5:**
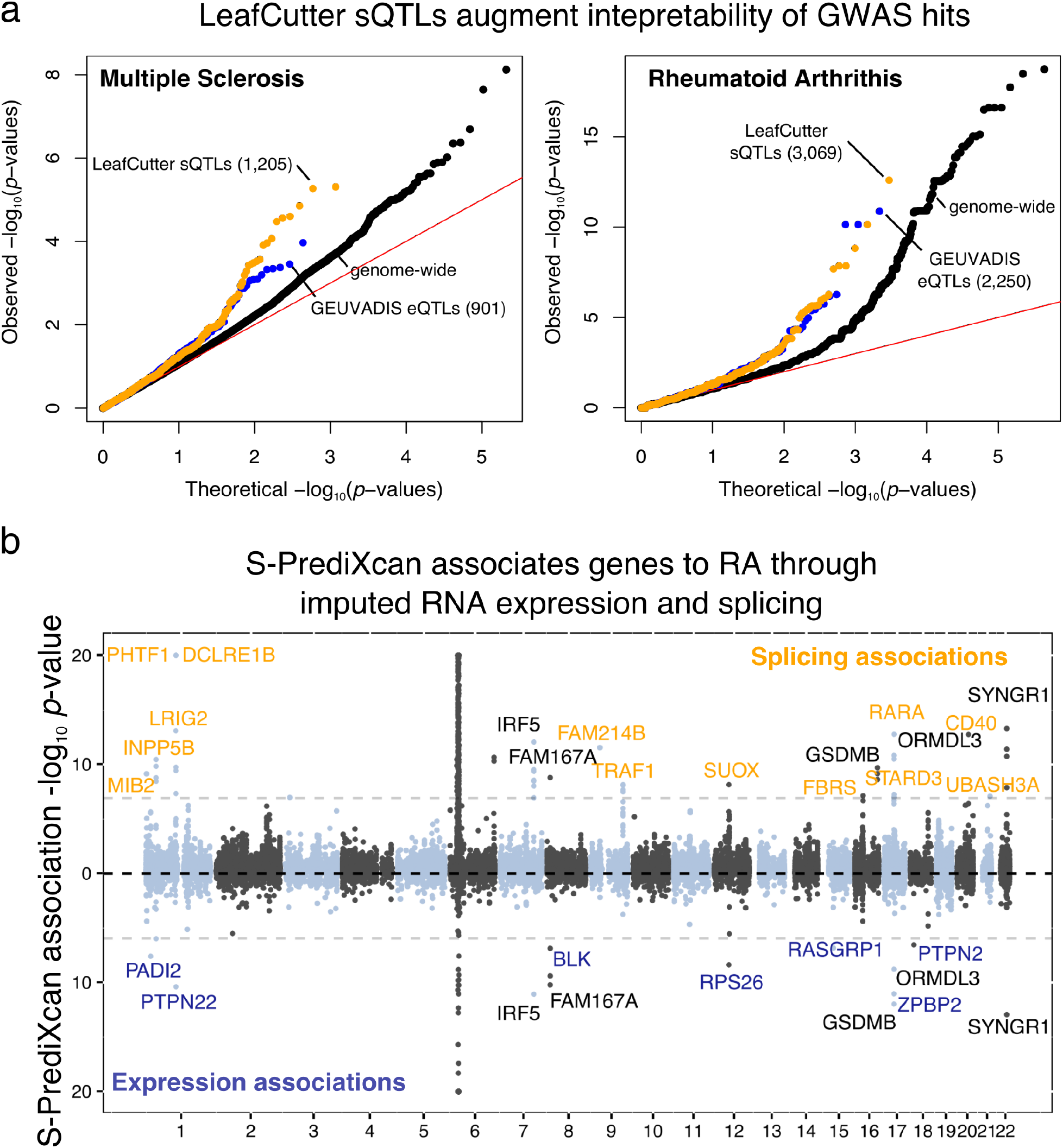
(a) Enrichment of low p-value associations to multiple sclerosis and rheumatoid arthritis among LeafCutter sQTL and GEUVADIS eQTL SNPs. The numbers of top sQTLs and eQTLs that are tested in each GWAS are shown in parentheses. (b) Manhattan plot of S-PrediXcan association p-values from prediction models for intron quantification (LeafCutter; top) and gene expression (GEUVADIS; bottom). Genes that were found to be associated through RNA splicing are highlighted in orange, those associated through gene expression in purple, and those associated through both in black. The names of associated genes from the extended MHC region are not shown.

To further explore the utility of LeafCutter sQTLs for understanding GWAS signals, we applied S-PrediXcan ^33^ to compute the association between predicted splicing quantification and 40 complex trait GWASs using models trained on GEUVADIS data (Methods and Supplementary Note 9). When applied to a rheumatoid arthritis (RA) GWAS, we found that considering intronic splicing allowed us to identify 18 putative disease genes (excluding genes in the extended MHC region), of which 13 were not associated using gene expression level measurements (Figure 5b). Novel putative disease genes associated through intronic splicing include CD40, a gene previously found to affect susceptibility to RA ^34^. However, we found no overall enrichment of functional categories among the 18 or 13 putative disease genes. Overall, using LeafCutter splicing quantifications allowed us to increase the number of putative disease genes by an average of 2.1-fold as compared to using gene expression alone (Supplementary Table 1). These results demonstrate that by dramatically increasing the number of detected sQTLs, LeafCutter significantly enhances our ability to predict the molecular effects of disease-associated variants.

In conclusion, our analyses show that LeafCutter is a powerful approach to study variation in alternative splicing. By focusing on intron removal rather than exon inclusion rates, we can accurately measure the stepwise intron-excision process orchestrated by the splicing machinery. Our count based statistical modeling, accounting for overdispersion, allows identification of robust variation in intron excision across conditions. Most importantly, LeafCutter allows the discovery of far more sQTLs than other contemporary methods, which improves our interpretation of disease-associated variants.

## Methods

### Identifying alternatively excised introns

To identify clusters of alternatively excised intron, split-reads that map with minimum 6nt into each exon are extracted from aligned .bam files. Overlapping introns defined by split-reads are then grouped together. For each of these groups LeafCutter constructs a graph where nodes are introns and edges represent shared splice junctions between two introns. The connected components of this graph define intron clusters. Singleton nodes (introns) are discarded. For each intron cluster, LeafCutter iteratively (1) removes introns that are supported by fewer than a number of (default 30) reads across all samples or fewer than a proportion (default 0.1%) of the total number of intronic read counts for the entire cluster, and (2) re-clustered introns according to the procedure above.

### Dirichlet-multinomial generalized linear model

Intron clusters identified from LeafCutter comprise of two or more introns. More specifically, each intron clusters C identified using LeafCutter consists of *J* possible introns, which have counts y*_ij_* for sample *i* and intron *j* (and cluster total n*_iC_* = Σ*_j'_* y_*ij*'_), and *N* covariate column vectors x*_i_* of length *P*. LeafCutter uses a Dirichlet-Multinomial (*DM*) generalized linear model (GLM) to test for changes in intron usage across the entire cluster, instead of testing differential excision of each intron separately across conditions or genotypes.

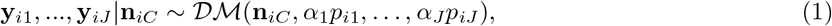

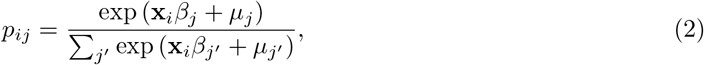

where (2) corresponds to the softmax transform, which ensures Σ*_j_ p_ij_* = 1. We perform maximum likelihood estimation for the outputs: the *J* coefficient row vectors *β_j_* of length *P*, the intercepts *µ_j_* and concentration parameters *α_j_*. We use the following regularization to stabilize the optimization:

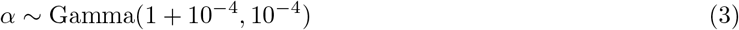

The Dirichlet-Multinomial likelihood is derived by integrating over a latent probability vector π in the hierarchy

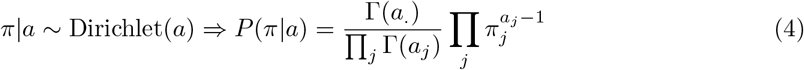

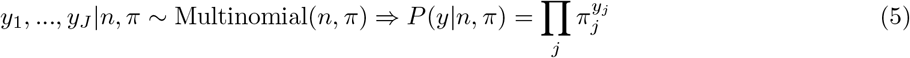

where a. = Σ*_j_ a_j_*, to give

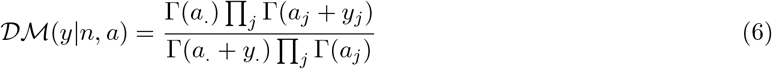

In the limit 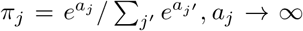 for all *j*, we have 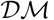 *(n,a)* → Multinomial(*n, π*). For the GLM this means that as α*_j_* → ∞ we recover a multinomial model with no overdispersion. Smaller values of α*_j_* correspond to more overdispersion.

While the Dirichlet-multinomial effectively accounts for overdispersion, it fails to handle extremely outlying in samples, which negatively impacts calibration. To reduce sensitivity to such outliers we developed a robust likelihood model

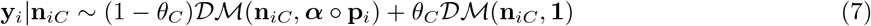

where *θ_C_* is a per-cluster mixture proportion giving the probability that a sample comes from the outlier distribution. Using 1 as the parameter vector for the outlier distribution corresponds to the underlying Dirichlet distribution being uniform over the simplex. θ_C_ is learnt jointly with the other parameters, and given a prior Beta(1.01, 10^-4^).

### Differential intron excision across conditions

To test differential intron excision between two groups of samples, we encode *x_i_* = 0 for one group and *x_i_* = 1 for the other in the Dirichlet-Multinomial generalized linear model. We apply two filters to ensure we only perform reasonable tests:

- Only introns which are detected (i.e. have at least one corresponding spliced read) in at least five samples are tested.
- A cluster is only tested if each group includes at least 4 individuals with 20 spliced reads supporting introns in the cluster.

The thresholds in these filters are easily customizable as optional parameters.

### Mapping splicing QTLs

To identify splicing QTLs, RNA-seq reads are mapped onto the genome using a RNA-aligner such as STAR ^35^ or OLego ^20^. Because LeafCutter only uses reads that map across junctions to estimate intron excision rates, it is essential to remove read-mapping biases caused by allele-specific reads. This is particularly significant when a variant is covered by reads that also span intron junctions as it can lead to spurious association between the variant and intron excision level estimates. Subsequent to mapping, LeafCutter finds alternatively excised intron clusters and quantifies intron excision levels in all samples. LeafCutter outputs intron excision proportions, which are used as input for standard QTL mapping tools such as MatrixEQTL or fastQTL (Supplementary Methods 6).

### S-PrediXcan analyses

Prediction models for intron quantification (LeafCutter) and gene expression (GEUVADIS) were trained using Elastic Net on GEUVADIS data. A value of α = 0.5 was chosen for the mixing parameter. Prediction performance for gene expression remains stable for a wide range of mixing parameters when α does not approach 0.0 (Ridge Regression) ^36,37^. For each gene, we used SNPs within 1Mb upstream of the TSS and 1Mb downstream of the TTS. Similar windows around each splicing clusters were chosen.

We downloaded a genome-wide association meta-analysis summary statistics for Rheumatoid Arthritis from http://plaza.umin.ac.jp/yokada/datasource/software.htm, and ran S-PrediXcan using these models. A total of 4,625 gene associations were obtained for the genetic expression model, and 41,196 intron quantification cluster associations for the splicing model, that had a model prediction False Discovery Ratio under 5%.

### Visualizing LeafCutter differential splicing output

Using the R Shiny framework and `ggplot2`, we created an interactive browser-based application, Leaf`Viz`, that allows users to visualize LeafCutter differential splicing analyses. Leaf`Viz` generates LeafCutter cluster plots with information on the significance of the detected differential splicing and the estimated differences of the splicing changes. All significant clusters are labelled as “annotated” or “cryptic” by intersecting junctions with a user-defined set of transcripts (e.g. gencode v19). Users can directly download plots from the website in PDF format, which can be easily edited for publication. An example of LeafViz applied to a differential splicing analysis between 10 brain and 10 heart samples from GTEx is available here: https://leafcutter.shinyapps.io/leafviz/.

## Acknowledgement

We thank Xun Lan and other members of the Pritchard Lab for helpful discussions and comments. This work was supported by a CEHG Fellowship, the Howard Hughes Medical Institute, and the US National Institutes of Health (NIH grants HG007036, HG008140, and R01MH107666).

## Author Contributions

Y.I.L., D.A.K. and J.K.P. conceived of the project. Y.I.L. and D.A.K. performed the analyses and implemented the software. D.A.K. developed and performed the statistical tests and modeling. J.H. implemented the visualization application. A.N.B., S.P.D., and H.K.I. performed the S-PrediXcan analyses. Y.I.L. and J.K.P. wrote the manuscript.

## Supplementary Material for Annotation-free quantification of intron splicing for genomic studies

Yang I Li1;^9^, David A Knowles^1;2;3;9^, Jack Humphrey^4;5^, Alvaro N. Barbeira^6^, Scott P. Dickinson^6^, Hae Kyung Im^6^, Jonathan K Pritchard^1;7;8^

## 1 Identifying alternatively spliced introns using LeafCutter

Starting from alignment files in .bam format, junctions from split-reads that map with minimum 6nt into each exon are extracted using a script we provide (1) based on two OLego helper scripts. Then, the LeafCutter clustering program (2) can be used to identify intron clusters supported by at least 30 (option `-m`) total reads (across all samples) and introns supported by more than 0.1% (option `-p`) of the total read counts for the entire cluster. The number of reads supporting each intron and cluster is then counted in all samples separately and collated in a table for downstream analyses.

**Figure S1:**
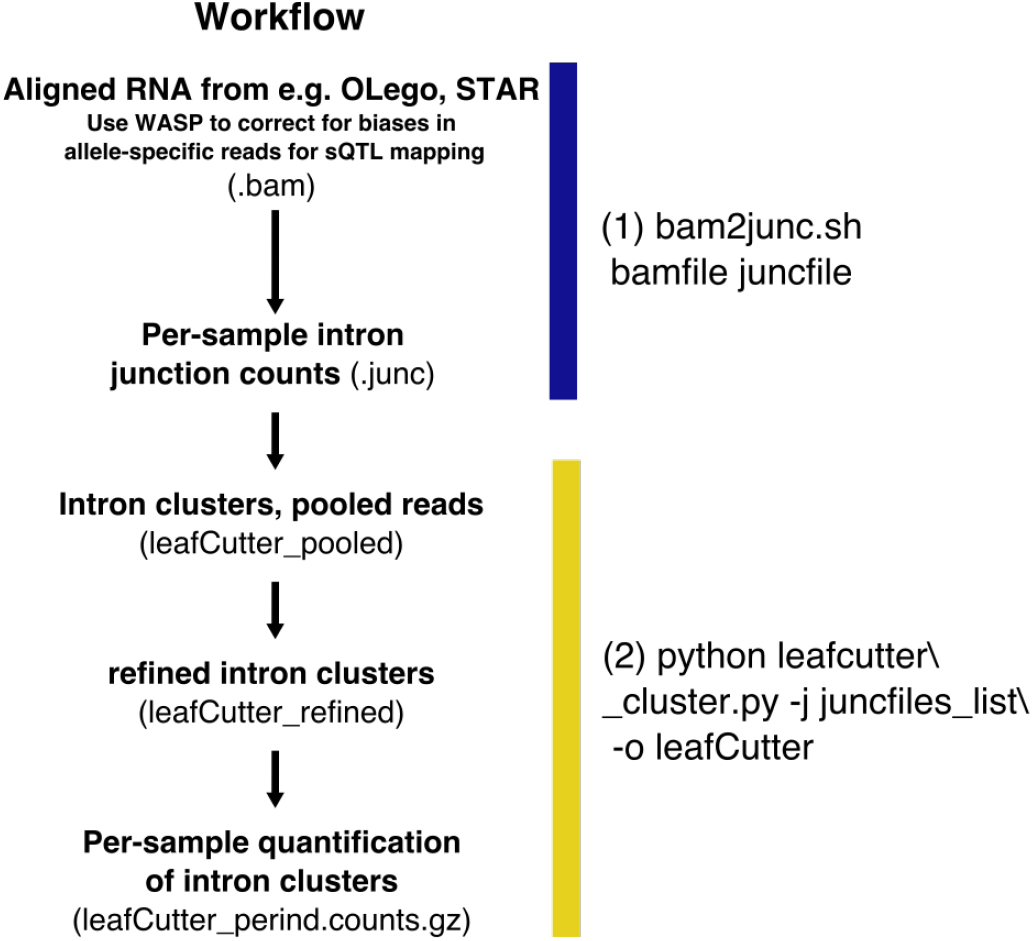
Helper method and LeafCutter workflow for intron clustering.

Because LeafCutter focuses on intron splicing rather than whole isoform quantification, alternative transcription start site or polyadenylation sites are not captured. However, several prevalent types of alternative splicing (Figure S2) are equivalent to specific intron excision events.

**Figure S2:**
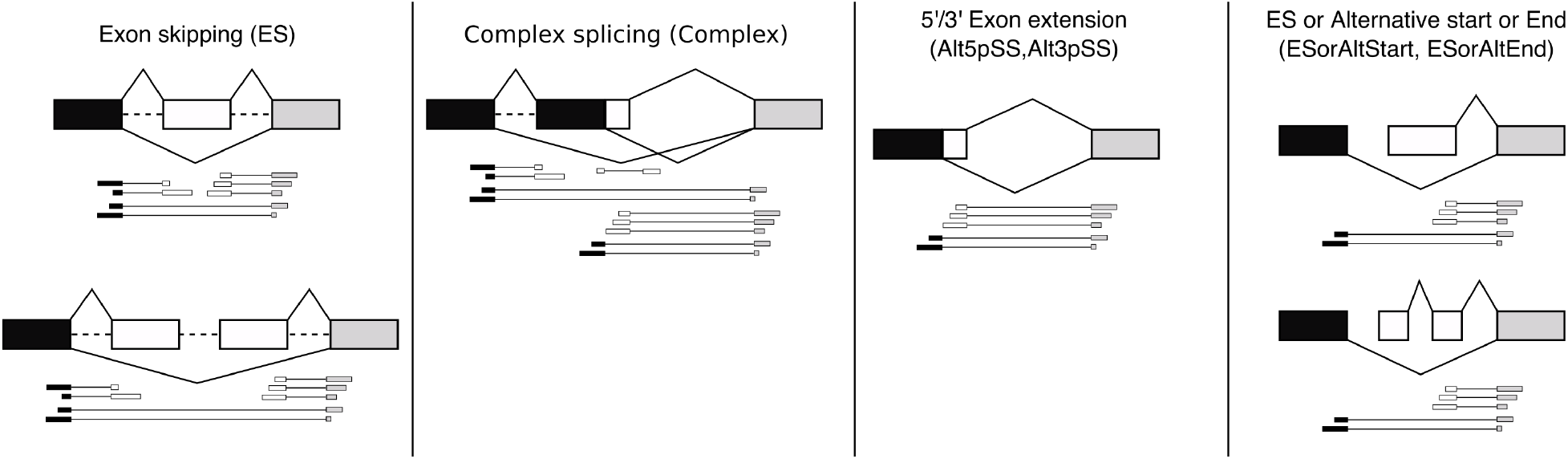
Several types of common alternatively splicing events are captured by the alternative excision of introns.

## 2 Comparison of LeafCutter to other methods for differential splicing analysis

### 2.1 rMATS, MAJIQ, and Cufflinks2

To compare the ability of different software to detect differential splicing, a fair comparison needs to overcome (1) differences in p-value calibration, and (2) differences in what is being measured e.g. transcript ratios versus local splicing events. As test data, we chose to contrast lymphoblastoid cell lines (LCLs) derived from Yoruba individuals against LCLs derived from central european (CEU) individuals. We chose LCLs as they are homogeneous cell lines and splicing differences between populations should be subtle; both properties are favorable for comparing sensitivity of the methods.

To overcome (1) the problem of p-value calibration, we computed the empirical false discovery rate (FDR) as follows:

a. First, we identify differential splicing between YRI and CEU LCLs using each method and record the p-value (1-posterior for MAJIQ, see subsection below) distribution for all tests.
b. Next, we permuted labels on the samples such that ∼ 1/2 of CEU samples are labeled as YRI samples and vice versa. We then run each method on these permuted samples and the p-value (1-posterior for MAJIQ) distribution are once again recorded.
c. The number of differential splicing events discovered at a certain FDR (e.g. 5%) is defined as the maximal number of events with test *p*-value less than *p* in the real data (*N_real_*) such that the number of events with test *p*-value less than p in the permuted data (*N_perm_*) respects the following constraints *N_perm_*/(*N_perm_* + *N_real_*) < *FDR*.

The resulting p-value distributions of the 3v3, 5v5, 10v10, and 15v15 comparisons are shown in (Figure S3). We observed that LeafCutter p-values were generally well-calibrated, which resulted in the largest number of differentially spliced events compared to rMATS, MAJIQ, and Cufflinks2.

We observed that Cufflinks2 p-values were very conservative (see Cufflinks subsection below). We therefore report the number of significantly differentially spliced events from Cufflinks2 directly. Interestingly, Cufflinks2 reports 19 significantly different splicing events in the 3v3 comparison, but not in comparisons with large sample sizes.

To overcome (2) the problem of differences in what events are being measured, we collapsed all events in rMATS and MAJIQ that shared a single splice site into a single event (as is done in LeafCutter).

### 2.2 MAJIQ

Instead of computing p-values for differentially splicing tests, MAJIQ computes posterior values reflecting the confidence that a splicing event is differentially spliced at a ∆Ψ of at least *P* which is an user-defined parameter. In our tests, we chose *P* to be 0.05. Choosing other values of *P*, e.g. 0.01 resulted in similar or worse performance.

In principle it should be possible to use the posterior probabilities from MAJIQ’s Bayesian model to directly control FDR. In particular, taking events with posterior probabilities ¿1-F should control FDR at F. However, our permutation analysis shows this is clearly not the case since this approach results in a highly inflated false positive rate (FPR) under the null. The fact that MAJIQ does not seem to give “true” posterior probabilities suggests some degree of model mis-specification, i.e. that the statistics of real RNA-seq counts do not quite match the assumptions made by the MAJIQ differential splicing model.

### 2.3 Cufflinks2

We sought to understand the source of Cuffdiff2’s overly conservative p-value distribution under the null. To test for differential isoform usage for a specific gene Cuffdiff2 considers estimated isoform usage proportions for a gene in two groups, denoted 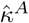 and 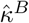 as well as associated posterior covariances 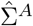 and 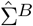. The test statistic used is the Jensen-Shannon distance (JSD), 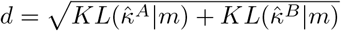 with 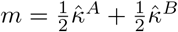. Under the null 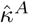 and 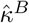 are drawn from the same distribution, which Cuffdiff2 assumes to be multivariate normal. To approximate the sampling distribution of d, 10^5^ *pairs* of samples are drawn from 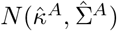, and the JSD for each pair. The procedure is repeated using 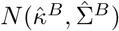 and the two resulting empirical *p*-values are averaged.

To test the calibration of this procedure we simulated an idealized scenario where 1000 reads in each of two conditions are unambiguously mapped to 5 isoforms of a gene. The true (shared) usage proportions are sampled uniformly from the 5-simplex. Per condition counts are sampled from a Dirichlet-multinomial distribution to model overdispersion, with a concentration parameter *c* = 10, typical for RNA-seq data. We obtained maximum likelihood estimates of 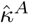 and 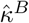 under the “best-case” scenario of knowing the true *c*, and corresponding 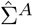 and 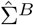 estimates using the inverse Hessian of the log likelihood function. We then performed the Cuffdiff2 procedure using these values. The whole procedure was repeated for 100 different simulated true usage proportions. This procedure recapitulates the overly conservative *p*-value distribution (Figure S4 and S3) we observed when applying Cuffdiff2 to permuted real RNA-seq data. We hypothesize that the root cause of the problem is that the multivariate normal is a poor approximation for distributions constrained to the simplex, and as a result the estimated sampling distribution of *d* is considerably more dispersed than it should be.

**Figure S3:**
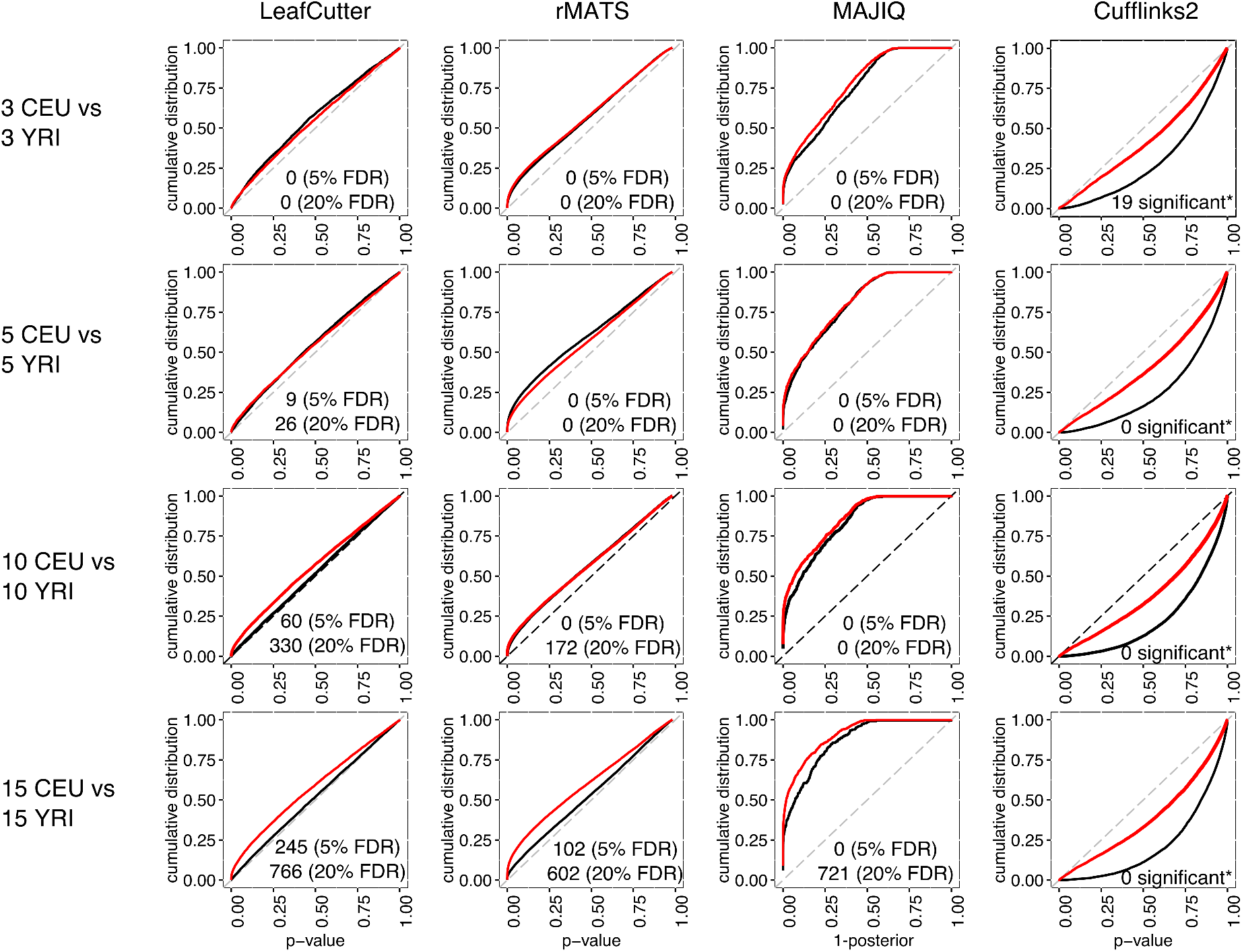
Cumulative distributions of differential splicing test p-values (1-posterior for MAJIQ) for the all YRI versus CEU LCLs comparison (red). The distribution of test p-values for the permuted comparisons are also shown (black). * Cufflinks2 reports 19 significantly differentially spliced genes in the 3 vs 3 comparison, but none in the other comparisons.

### 2.4 Comparison of false negative rates

To evaluate the false negative rates of differential splicing methods, we simulated sequencing reads for 160 protein coding genes each with 2 to 15 transcripts. For each gene, we only considered transcripts that differed by at least one overlapping intron when compared to another transcript to avoid cases where two transcripts only differ e.g. in the first or last exon or in an intron retention event (neither of which LeafCutter aims to detect). We then simulated reads from these transcript models as follows:

**Figure S4:**
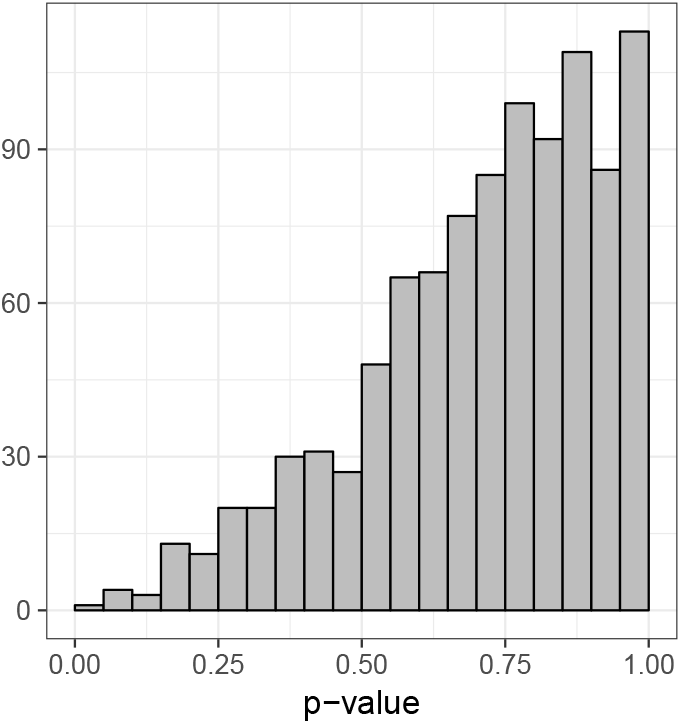
Simulated isoform usage under the null of no differential splicing shows Cufflinks2 p-values are overly conservative.

1. We simulated 8 biological samples each with 5 technical replicates.
2. For each gene, we set a random transcript’s expression to 1X (no change), 1X, 1X, 1.1X, 1.25X, 1.5X, 3X, and 5X in the 8 biological samples in random order (note that we set 1X for 3 of 8 samples, so there are 3 comparisons with no change of transcript expression; we used these to compute false positive rates).
3. We used polyester ^38^ to simulate sequencing reads, obtaining 8 × 5 = 40 RNA-seq samples (we used default parameters, e.g. 30X coverage and default error distributions).
4. We mapped reads from each sample using STAR and applied all four differential splicing detection methods on all pairwise (8 choose 2) = 28 comparisons.
5. We computed the effective transcript fold-change for each gene (a transcript might be set to 1.5X and 3X in the two samples that are being compared resulting in a effective fold change of 2X) in all 28 pairwise comparisons.
6. We then collected all p-values for every gene/comparison (min p-value/max posterior if more than one splicing event is tested per gene) and plotted their differential splicing test p-values binned by their effective transcript fold-change (Figure S5).
7. For each effective transcript fold-change, we computed the true positive and false positive rates for all possible p-value or posterior cutoffs (Figure S6).

From these simulations and the receiver operating characteristic (ROC) curves, we conclude that while Cufflinks2 appears to detect more transcripts with 1.1 fold-difference at reasonable FDRs, LeafCutter outperforms all three other methods when transcripts differed by 1.25-fold or more (Figure S6). Of the four methods tested, Cufflinks2 is the only method that estimates transcript levels, which might explain its higher power in detecting small differences in transcript expression. Interestingly, the performance of MAJIQ and LeafCutter were nearly identical when evaluated on transcripts that differed by 3-fold or more, but LeafCutter outperformed MAJIQ when differences were more subtle. This can be explained by the observation that LeafCutter has a lower false positive rate than compared to MAJIQ (see LeafCutter and MAJIQ panels at 1X effective fold-change in Figure S5).

### 2.5 Additional comparisons

As further comparisons and to ensure that the differentially spliced events detected using LeafCutter are not simply noise. We first asked about the correlation of p-values between comparisons with varying sample sizes. Here, we only compared LeafCutter to rMATS as MAJIQ do not report p-values and Cufflinks2 p-values are overly conservative. To do this, we computed the Spearman correlation of the – log *p* of the tested introns in the 15v15 comparison versus the corresponding – log *p* of the tested introns in the 3v3, 5v5 and 10v10 comparisons. As expected, the correlations increase monotonically for both methods as sample size increases reflecting an increase in precision in our effect size estimates (Figure S7a). However, we do observe a significantly higher correlation for LeafCutter compared to rMATS, suggesting that LeafCutter is more robust to comparisons involving fewer samples.

We further observed that the ability of LeafCutter to recall genes with evidence of differentially splicing discovered using an rMATS analysis was similar to that of MAJIQ, while Cufflinks2 showed the worst performance of all (Figure S7b).

To estimate the concordance between methods, we ranked genes by our differential splicing p-values, and asked about concordance at different bins of significance levels (50 genes per bin). We found that for the most significant bin (i.e. the top 50 most significantly differentially spliced genes), the concordance was high (65–75%) between rMATS and LeafCutter (we used a p-value cutoff of 0.05 to determine concordance) and even higher (80–82%) between LeafCutter or rMATS top genes and MAJIQ genes (we used a posterior > 0.99 to determine concordance). These observations (Figure S26a,b) are in line with our expectation that concordance rates rapidly decrease as our power to detect differentially spliced genes drops to zero.

Because LeafCutter, rMATS, and MAJIQ all measure splicing at a local level and not at the gene/isoform level, we next verified how consistent LeafCutter was with other predictions in terms of these local events. To this end, we ranked LeafCutter associations in terms of their p-values and asked whether LeafCutter introns shared at least one splice site with introns that were predicted to be differentially spliced by rMATS (*p* < 0.01) and MAJIQ (posterior > 0.95). We found that ∼ 90% of the introns that were found to be most significantly differentially spliced using LeafCutter shared a splice site with rMATS and MAJIQ, suggesting that LeafCutter identified the same differentially spliced events (Figure S26c). In contrast, only ∼ 60% of the events shared a spliced site when no associations was in LeafCutter (*p* > 0.5). Although 60% might appear high for the sharing between two “random” introns, it is useful to note that these are conditioned on introns that show (1) alternative splicing and (2) are differential spliced in rMATS or MAJIQ. The random overlap between LeafCutter-tested introns and rMATS-tested introns is less than 20%.

### 2.6 RAM usage

To measure RAM usage across methods, we used a custom script which calls strace `-e trace=mmap,munmap,brk` on the main programs, except for rMATS. We found that `rMATS` launched additional processes that are not measured directly. We therefore ran our custom script on `rMATS.3.2.5/processGTF.BAMs.py` which appears to be the most RAM intensive script of the rMATS pipeline.

**Figure S5:**
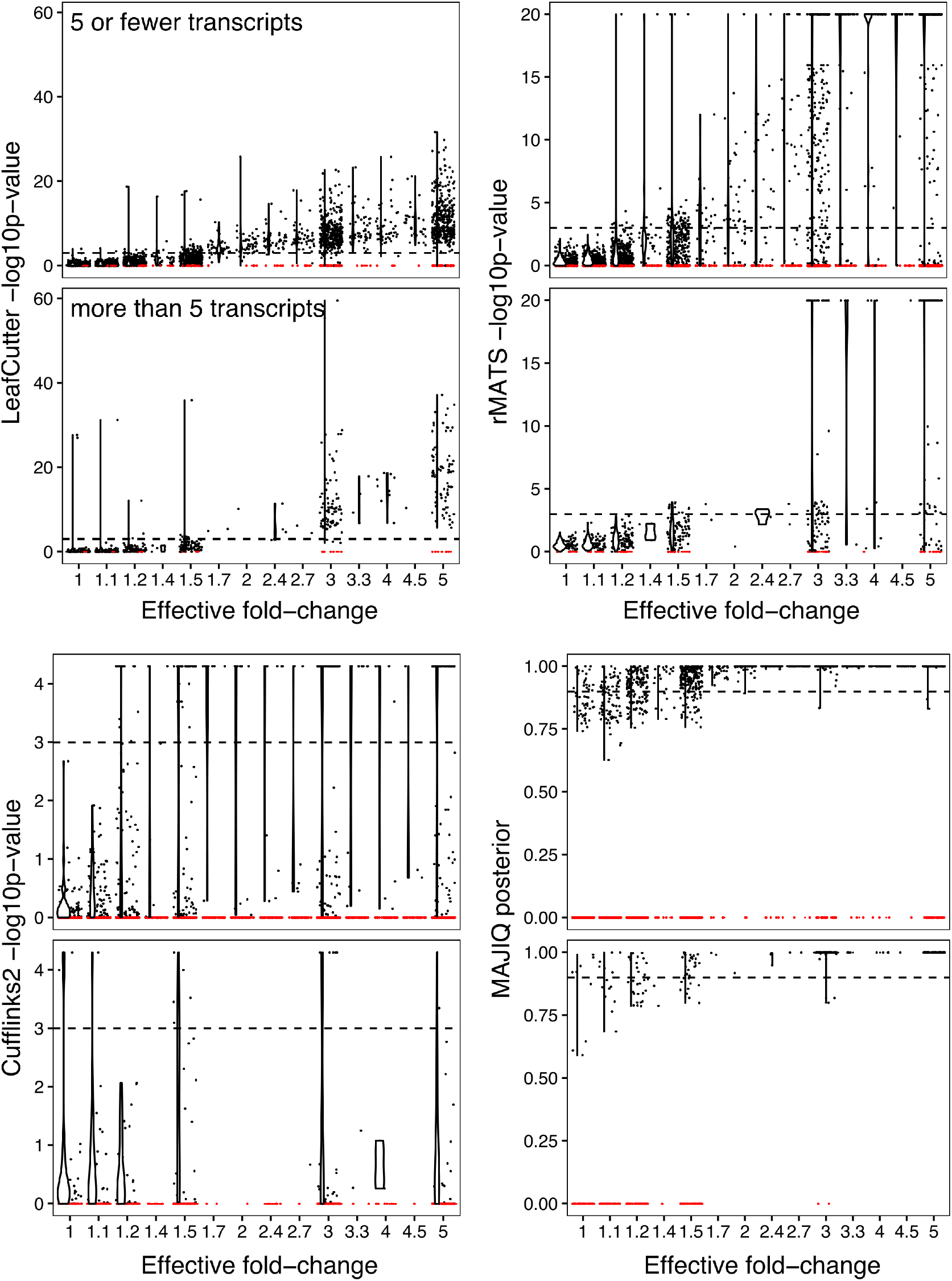
Scatter and violin plots of the p-value and posterior distribution of differential test statistics binned by true, simulated, effective transcript fold-change. For each method, tests of genes with ffve or fewer transcripts and genes with more thanffve transcripts are plotted on the upper and bottom panels, respectively. We observed a decrease in power to detect differential splicing as transcript number increases using Cufflinks2, but not for the three other methods. Red dots represent genes with no tested splicing event.

**Figure S6:**
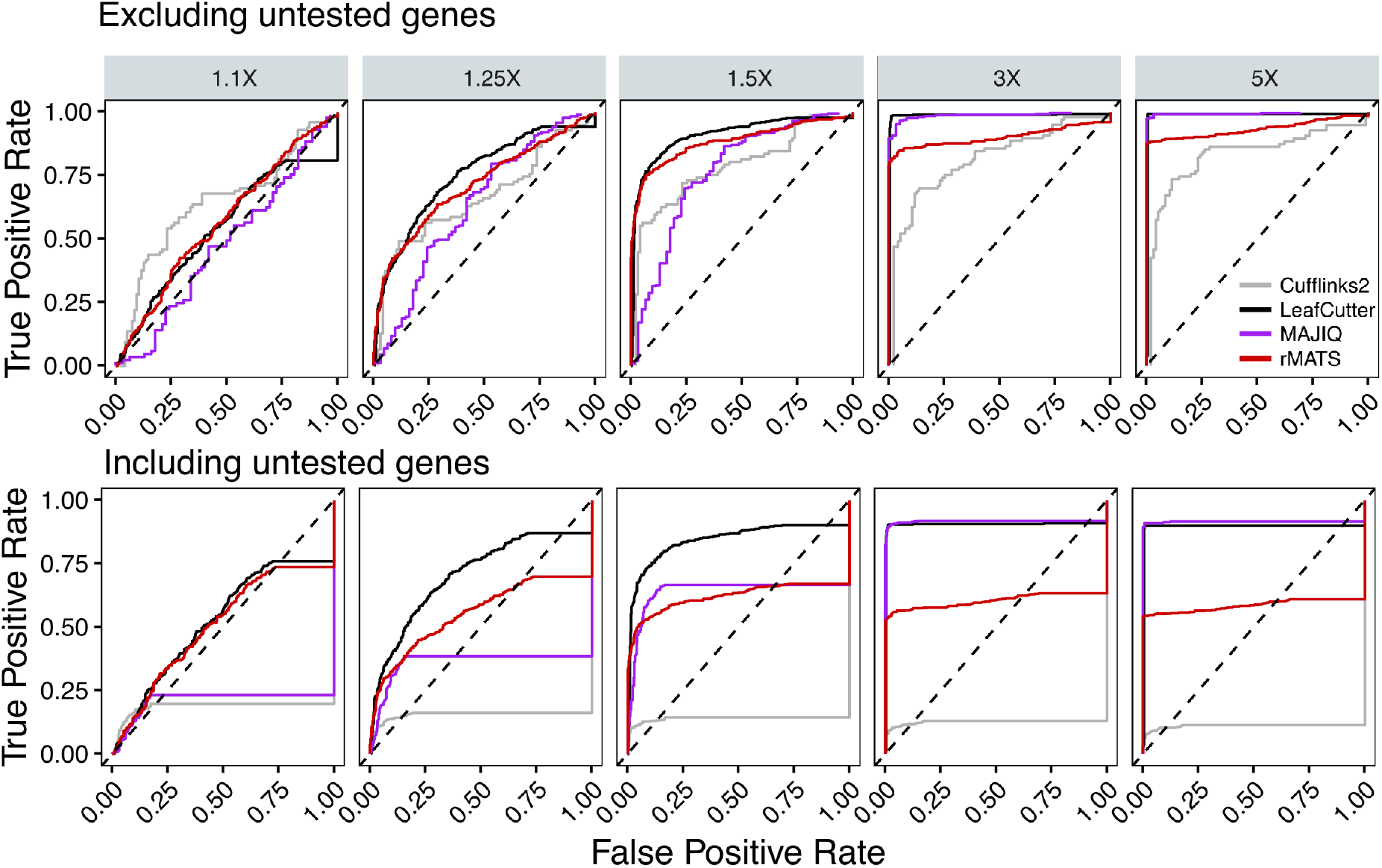
Receiver operating characteristic (ROC) curve of LeafCutter, Cuffinks2, rMATS and MAJIQ when evaluating differential splicing of genes with transcripts simulated to have varying levels of differential expression. Top panel shows ROC curves when excluding genes that were not tested by each respective methods. While the bottom plot includes genes that were not tested in the calculation of true positive rate.

**Figure S7:**
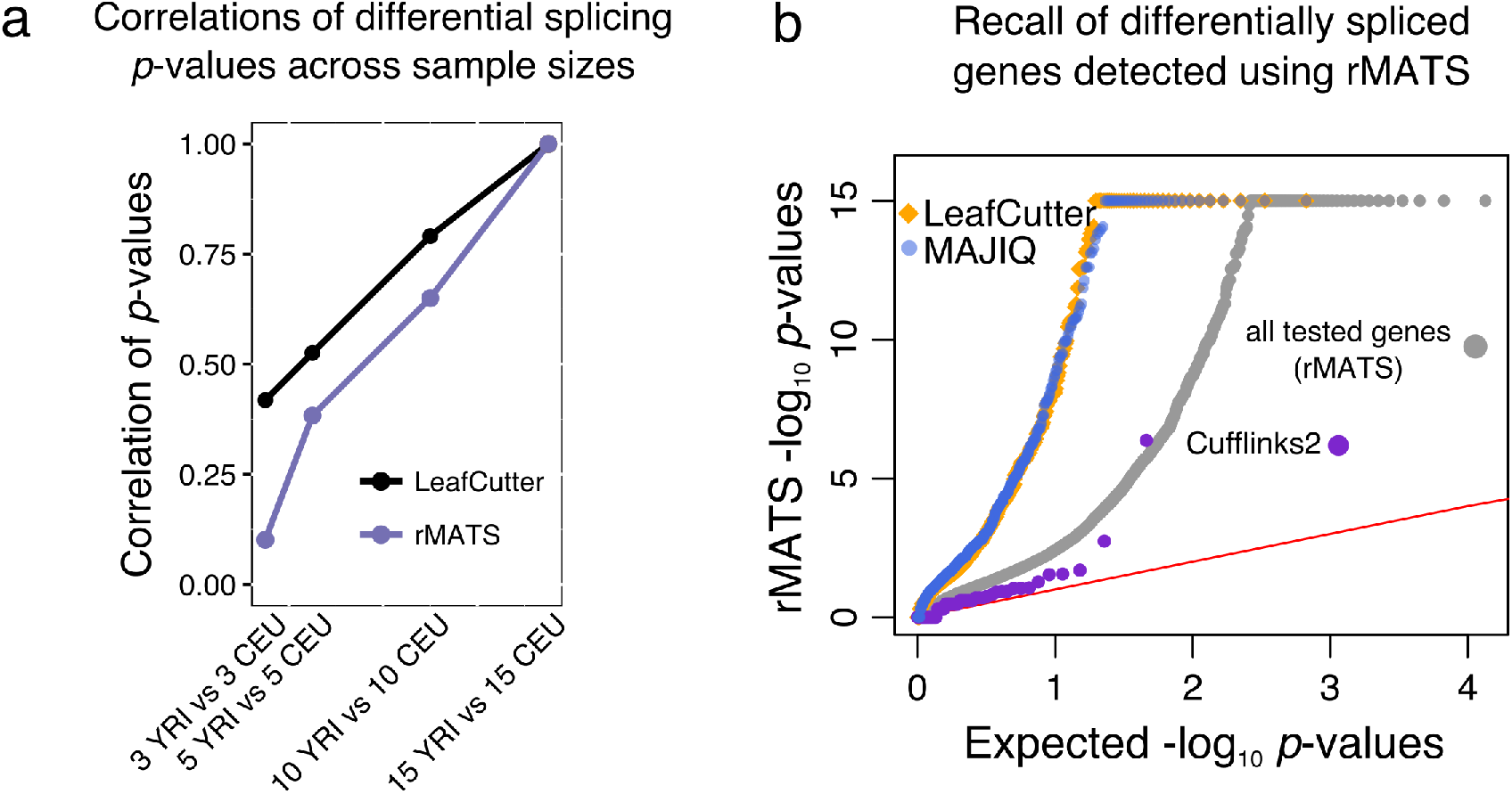
(a) Correlation of computed differential splicing – log_10_ (*p*-values) of introns between a 15 YRI vs 15 CEU LCLs comparison and 3 vs 3, 5 vs 5, and 10 vs 10 comparisons. (b) QQ-plot of the differentially splicing signal found using rMATS in a comparison between 15 YRI and 15 CEU LCLs samples. Differentially spliced genes detected using LeafCutter and MAJIQ, but not Cufflinks2, are highly enriched in genes detected using rMATS.

**Figure S8:**
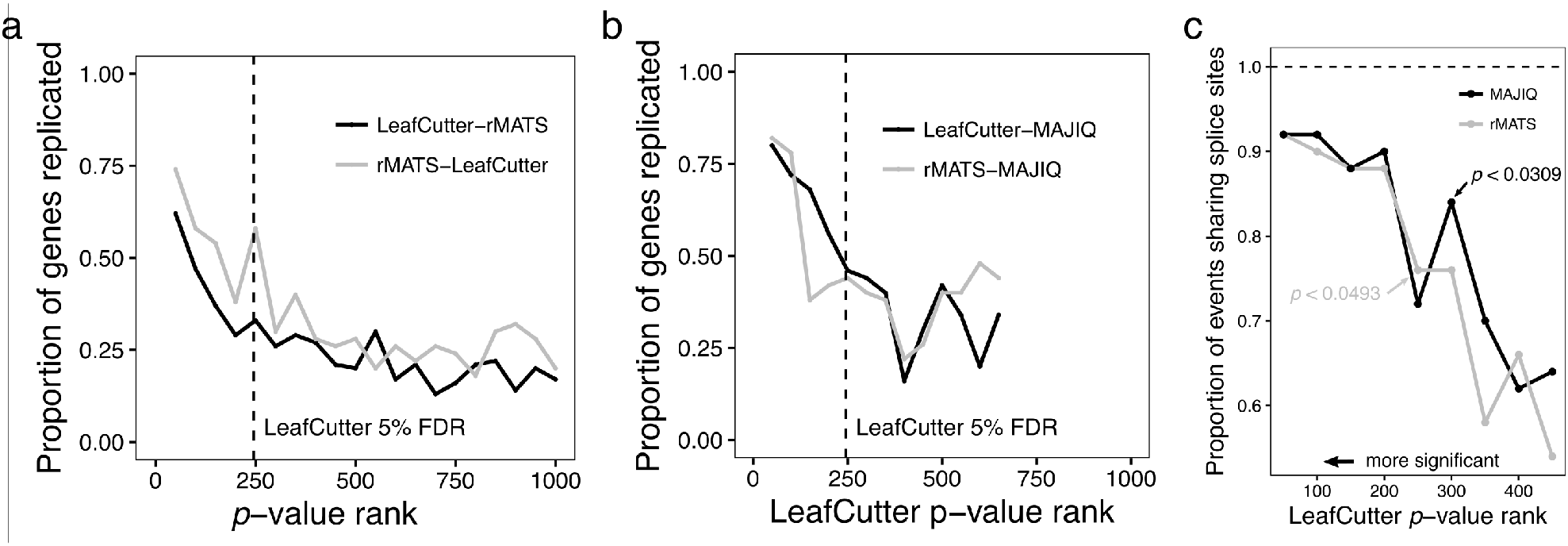
Estimates of concordances between differentially spliced genes detected using LeafCutter and rMATS genes (a) and between LeafCutter or rMATS genes and MAJIQ genes (b). Genes were ranked in terms of their signiffcance levels (from LeafCutter and rMATS) and grouped into bins of size 50. Dashed lines mark 245, i.e. the number of differentially spliced genes detected using LeafCutter at 5% FDR. (c) Estimates of the proportion of shared splice sites between differentially spliced introns predicted using LeafCutter and introns predicted to be differentially spliced using rMATS and MAJIQ. Genes were ranked in terms of their signiffcance levels (LeafCutter) and grouped into bins of size 50.

**Figure S9:**
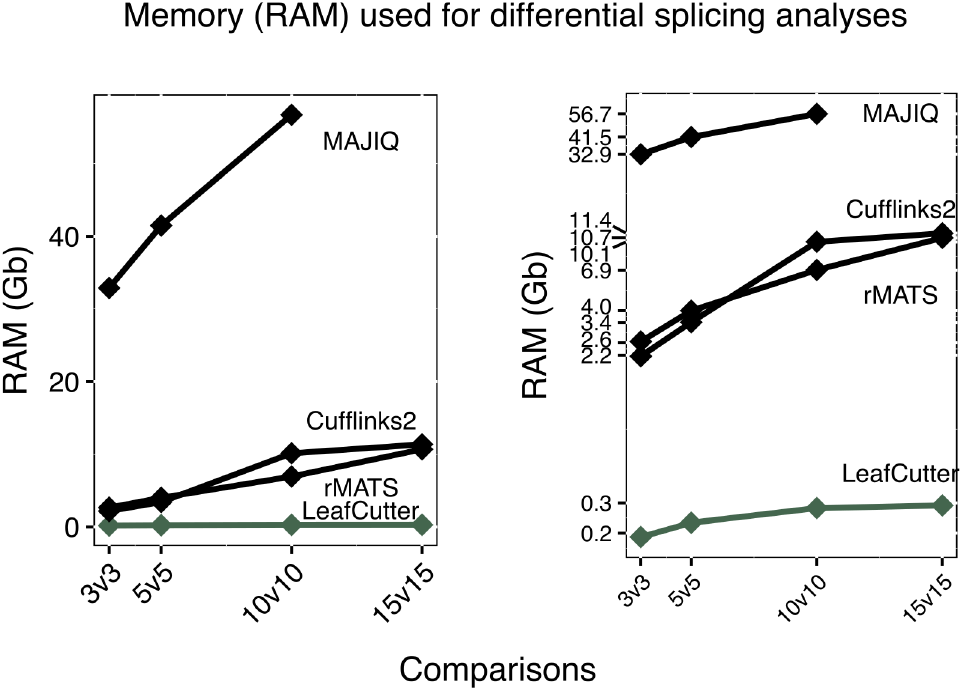
Memory usage (RAM) of four differential splicing methods applied to comparisons between 3, 5, 10, and 15 YRI vs CEU LCLs RNA-seq samples. We omitted the 15v15 MAJIQ run due to its expensive resource usage (both in terms of time and RAM). Right panel shows usage in log scale.

## 3 RNA-seq data processing

### 3.1 GTEx for intron discovery

We downloaded 2,192 RNA-seq samples from GTEx (Table S9). To analyze these, we used OLego (v1.1.5) ^20^ to map the RNA-seq reads to the human genome (hg19) and processed the resulting .bam files using Leaf-Cutter. Specifically, we used the following command: `olego -v -j hg19.intron.hmr.brainmicro.bed -e 6 hg19.fa`.

**Table S9:** Sample sizes of processed GTEx RNA-seq short read data by tissue type.

| Tissue | Sample Number |
| --- | --- |
| Heart | 153 |
| Testis | 67 |
| Spleen | 7 |
| Skin | 340 |
| Brain | 422 |
| Colon | 86 |
| Blood | 270 |
| Pancreas | 66 |
| Adipose Tissue | 172 |
| Lung | 151 |
| Esophagus | 238 |
| Muscle | 176 |
| Kidney | 8 |
| Liver | 35 |

The choice of OLego ^20^ is based on our previous experience that it performs well for discovering unannotated exons of small length (e.g. 9nt micro-exons) ^39^. OLego is a program specifically designed for de novo spliced mapping of mRNA-seq reads, while STAR ^35^ does best when a set of junction is provided. Since a chief objective of our GTEx analysis was to identify novel exons and to identify conserved alternative splicing events across multiple species with annotations worse than that of human, we used OLego for our GTEx differential splicing analyses (we used STAR for sQTL analyses because of fast running time and high accuracy in mapping). To quantify the differences in mapping of the two aligners, we picked at random five RNA-seq samples from the GTEx consortium that we previously aligned using OLego and re-aligned them using STAR. We next analyzed the correspondence between the number of junction reads for each junction across the two aligners. We found that while there are junctions whose read counts are orders of magnitude different, only 4.8% of junctions differed by a count fold-difference of 1.1 or more (0.94% of junctions differed by a count fold-difference of 2 or more).

**Figure S10:**
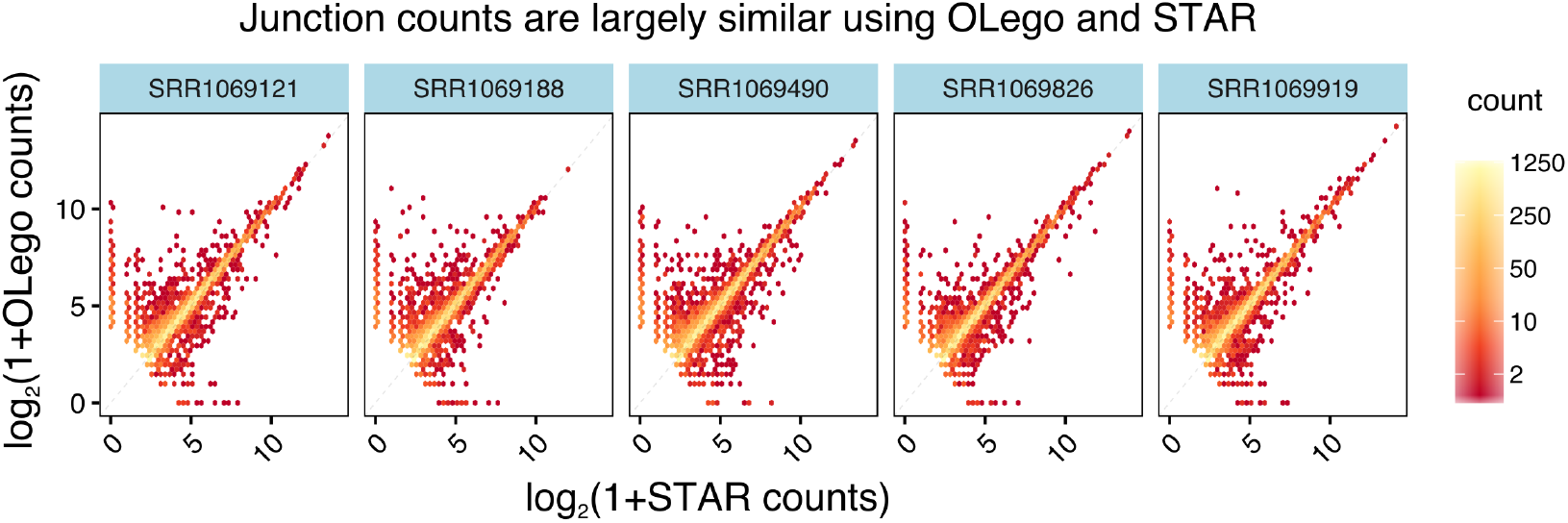
Comparisons of junction read numbers between STAR and OLego across 5 random GTEx samples. Only junctions with total reads of more than 16, across both aligners, are shown. Note that only junctions which were found using OLego in a bigger panel of GTEx tissues (i.e. all GTEx samples in this study) were considered.

### 3.2 GEUVADIS (YRI) for sQTL methods comparison

To compare splicing QTL (sQTL) calling methods, we aligned 85 YRI LCL samples from GEUVADIS using STAR two-pass and used WASP to remove reads that mapped with allelic biases ^18^. These aligned reads were used as starting point for each of the sQTL calling methods. Specifically, the following command was used:

`STAR ––genomeDir STAR_index ––twopassMode ––outSAMstrandField intronMotif ––readFilesCommand zcat ––outSAMtype BAM Unsorted`

### 3.3 GEUVADIS (CEU) for sQTL mapping

To control for differences in mapping procedures, we downloaded the .bam files directly from ArrayExpress (E-GEUV-3) and processed them using LeafCutter to obtain intron clusters and quantifications. We recommend the use of WASP ^18^ to correct for biases caused by allelic reads. However, to make our comparison to other tools fair, we used the aligned reads available on ArrayExpress, and removed all clusters with an association to a SNP that overlap junction reads (see section entitled “sQTL mapping using LeafCutter”). This approach is conservative as some allelic reads do not map with a bias.

### 3.4 GTEx for sQTLs mapping

Again, to control for differences in mapping procedures, we used the .bam files provided by the GTEx consortium for sQTL mapping, and removed all clusters with an association to a SNP that overlap junction reads.

## 4 Identification of unannotated introns in tissues from GTEx

To obtain a comprehensive set of annotated introns, we downloaded the GENCODE (v19), UCSC, and RefSeq annotation databases in .gtf format. We classified introns as annotated if their 5’ and 3’ splice sites correspond to the end and start, respectively, of two consecutive exons in at least one transcript. As such it is possible that both 5’ and 3’ splices sites of a novel intron are annotated. We note that although a large proportion of annotated introns are present in all three databases, we found that the GENCODE annotation has the most comprehensive list of introns.

To estimate the number of unannotated alternatively excised (AE) introns, we first mapped 2,192 RNA-seq samples from 14 tissues (GTEx) to the human genome (hg19) using OLego, allowing *de novo* splice junction predictions. We then used LeafCutter to identify alternatively excised introns by pooling all junction reads. We then restricted our analyses to AE introns that were supported by at least 20% of the total number of reads that support introns from the clusters they belong to in at least 25% of all samples, considering each tissue separately. Although there is no minimum read count (an intron supported by 20 reads, 20% of 100, is less likely to be the outcome of noisy splicing than one supported by 2 reads out of 10), we reasoned that requiring 20% percent-splicing in 25% of all samples will filter out most sequencing technical artifact and noisy splicing. Importantly, using different cutoffs does not alter qualitatively our conclusions. This resulted in 70,722 AE introns that met these criteria, of which 22,278 (31.5%) AE introns were absent from all three annotation databases.

To investigate the functionality of these unannotated introns, we asked whether the unannotated splice sites of the 22,278 AE novel introns show signature of sequence conservation across vertebrates. To do this, we divided splice sites into three classes: (1) control splice sites, which are annotated in one or more databases, but whose cognate splice site is unannotated, (2) the cognate splice site itself, and (3) splice sites of introns, for which both splice sites are unannotated. To compute sequence conservation, we average the phastCons score of the predicted splice sites (over 96% of which are AG/GT) plus 2 flanking bases. Interestingly, we find that the average sequence conservation of unannotated splice sites is higher if its cognate splice site is annotated (Figure 1d, Figure S11).

### 4.1 Validation of unannotated junctions in Intropolis

To verify that these unannotated splicing events are not a result of mapping errors or artefact unique to samples from GTEx, we examined the number of splicing junctions that could also be found in the Short Read Archive (SRA) using Intropolis ^24^ (note that GTEx samples were excluded from the SRA). Intropolis processed 21,504 human RNA-seq samples from the Sequence Read Archive (SRA) using RAIL-RNA to align and refine junction calls to improve sensitivity ^40^. This analysis therefore provides an additional replication of the RNA-seq aligner (i.e. OLego) that we used to identify unannotated splicing events. Because the SRA does not collect uniform cell-type or tissue labels for each sample, we used the cell-type or tissue labels predicted by phenopredict ^25^ to assign tissue identity to each SRA sample. Using this data, we quantified the number of alternatively spliced junctions identified in our study that can also be found in SRA samples (Figure S12). Overall, we found that, for instance, 86% of all novel junctions identified in GTEx testis using LeafCutter could be replicated in testis samples from the SRA (94% of unannotated heart junctions could be found in heart SRA samples). This is particularly impressive because (1) at most 56% of all unannotated junctions could be found in any other SRA tissues and (2) considering all tissues together increased the proportion of unannotated junctions “replicated” by only 4%, to 90%. These observations cannot be simply explained by a better sampling of testis in the SRA, as, for example, only 77% of the novel heart junctions could be found in SRA testis samples versus 94% in SRA heart samples.

**Figure S11:**
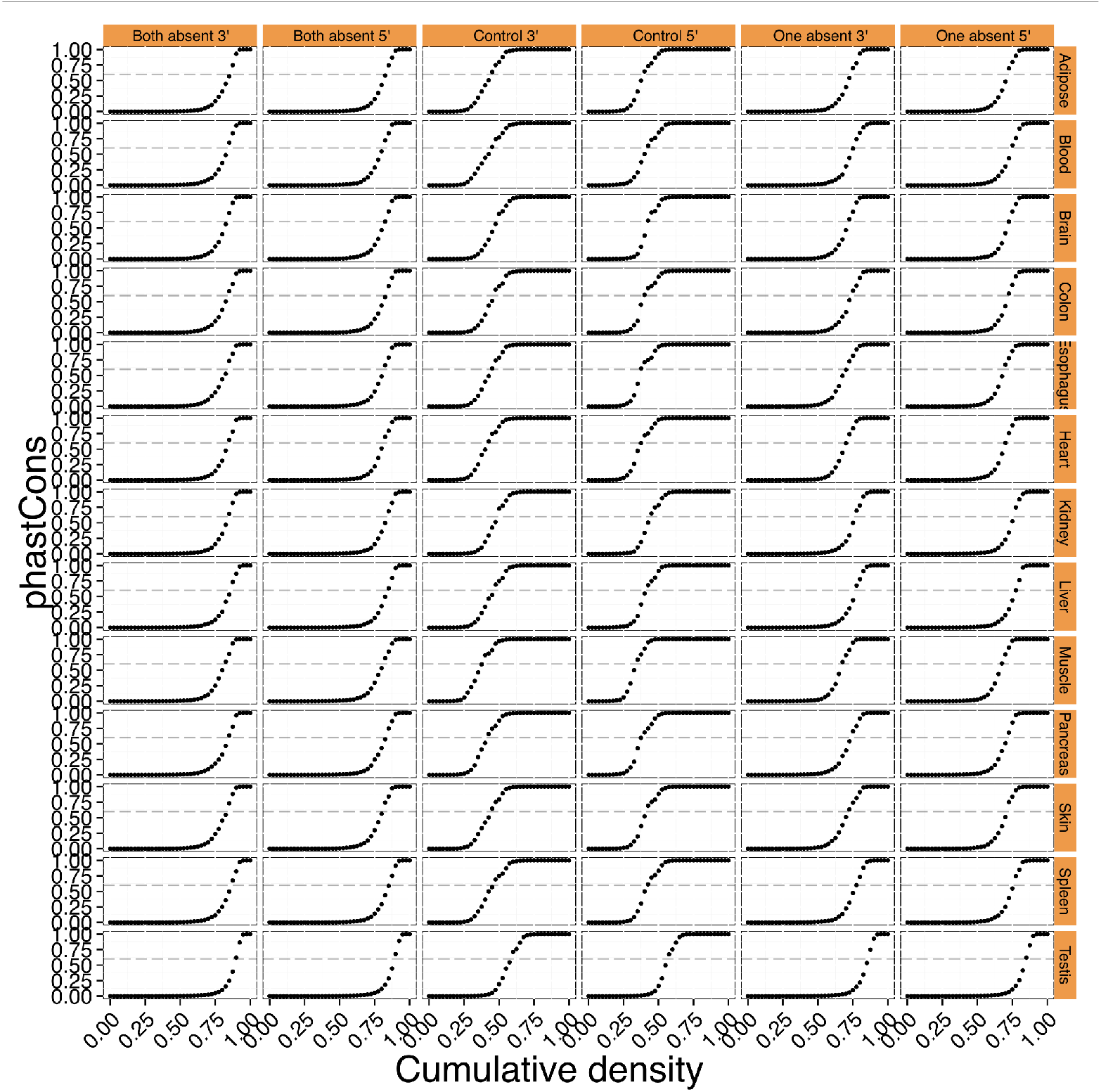
PhastCons score distribution of splice site of novel introns. While ∼60% of annotated splice sites have local phastCons score >0.6, only 15-25% of unannotated splice sites do. Thus ∼80% of novel splice sites may represent noisy intron excision events.

**Figure S12:**
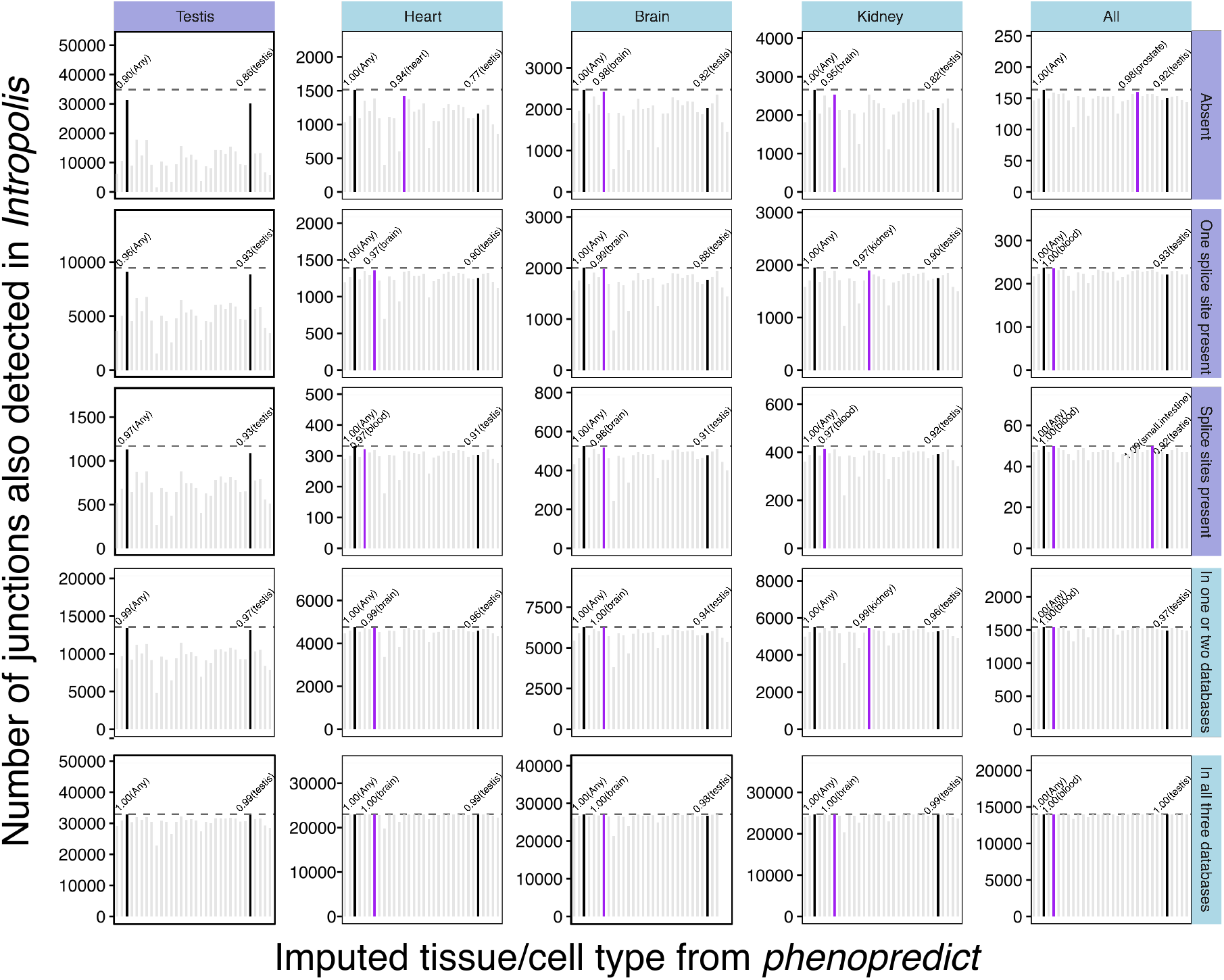
Barplots showing the number of alternatively used junctions annotated from our GTEx analyses that were found in Intropolis ^24^. phenopredictwas used to predict the tissue type corresponding to the SRA samples analyzed in Intropolis. For each set of junctions, the proportion of junctions that were found (at least 1 read) in any SRA sample (Any), or found in samples which were predicted to be from testis (Testis) are highlighted. The predicted tissues with the highest number of supported junctions are colored in purple. Eighty-six percent of all novel alternatively used testis junctions from our LeafCutter analysis could be found in testis samples within SRA (not including GTEx).

Because the analysis above only quantified presence or absence of the unannotated junctions in at least one sample from each tissue, we next characterized unannotated junctions by examining the proportion of samples in which they could be found by tissue (Figure S13). As expected, we found that unannotated junctions discovered in a given tissue tend to be present in a significant higher proportion of samples from the same, corresponding, tissue. Again, this suggests that unannotated junctions likely represent real splicing events that were not previously annotated as they tend to be highly tissue-specific.

To further profile this set of unannotated introns, we quantified their tissue-specificity, their levels of usage, and the type of splicing patterns they generally correspond to. As expected, we found that the vast majority of novel junctions were present in only a single GTEx tissue (Figure S14a). Similarly, we found that novel junctions identified in a tissue were used at a significantly higher levels in the corresponding tissue than in other tissues (Figure S14b), the differences were particularly striking for novel junctions discovered in testis. When we characterized the type of splicing events in which the unannotated introns were apart of, we found that, interestingly, 31.7% of all clusters with unannotated introns were complex, i.e. included at least one exon skipping and one alternative splice site event. This is nearly twice as many as compared to the 16.6% of complex clusters that are annotated. Overall, we conclude that unannotated junctions are relatively lowly used, tend to be tissue-specific, and often involve complex splicing patterns.

**Figure S13:**
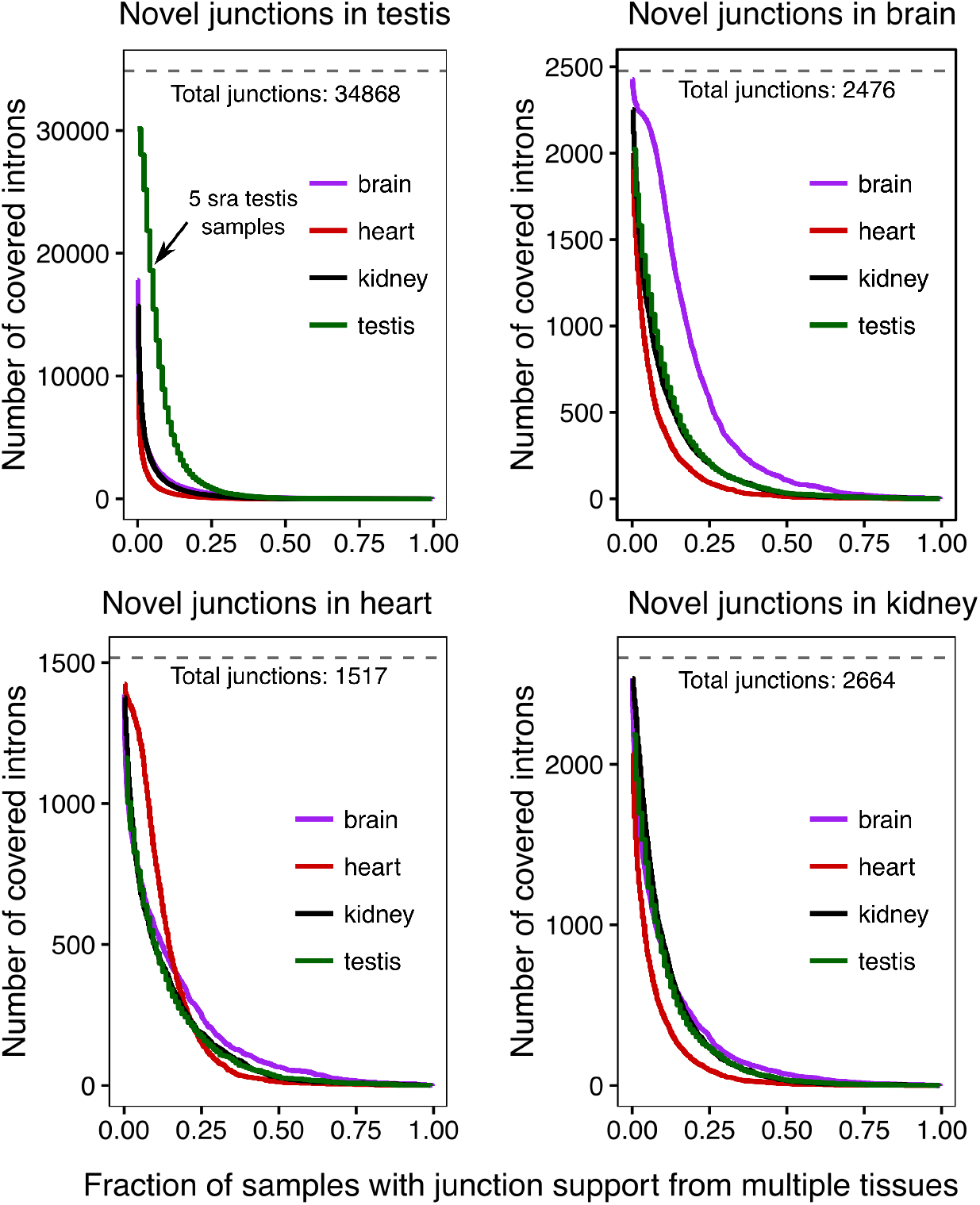
Number of junctions that were found in at least X percent of all SRA samples, by tissue.

**Figure S14:**
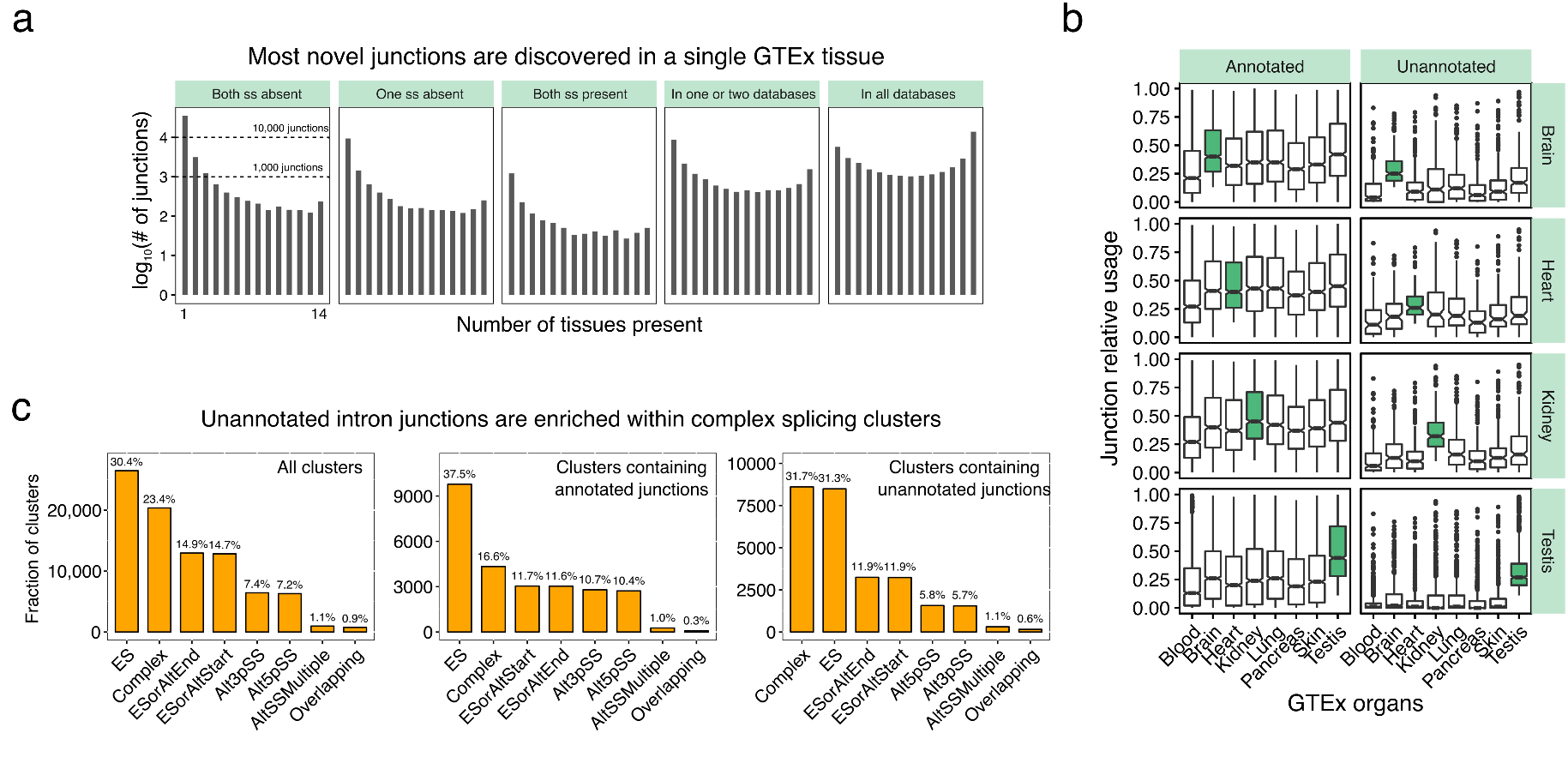
(a) Distribution of the number of different GTEx tissues in which junctions predicted to be absent, or present in three commonly-used annotation databases, could be detected. (b) Relative junction usage in multiple GTEx organs of annotated and unannotated junctions identified in four GTEx organs. (c) Distribution of LeafCutter clusters from GTEx samples in terms of their splicing types. Clusters with only annotated junctions and clusters with unannotated junctions were further separated.

## 5 Statistical models

For cluster C containing *J* possible introns, let *y_ij_* denote the count for sample *i* and intron *j* (and cluster total *n_iC_* = Σ*_j'_ y_ij'_*), and xi denote a *P*-vector of covariates.

### 5.1 Beta-binomial GLM

Our initial approach was to test each specific intron *j* of a cluster using

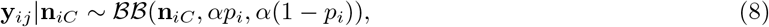

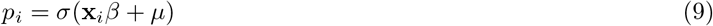

where *BB* is the beta-binomial distribution and *σ (x)* = 1/(1 + *e^-x^*) is the logistic function. Here the parameters to be learnt are the *P*-vector *β*, intercept *µ* and concentration parameter *α*. Higher values of *α* correspond to the underlying beta distribution concentrating around pi, and therefore to less count overdispersion. In particular as *α →* ∞ the *BB* likelihood converges to a multinomial likelihood, recovering a logistic regression model.

**Optimization**. For both the beta-binomial and Dirichlet-multinomial models we use the Bayesian probabilistic programming language Stan ^41^ to define the model, generate efficient C++ code for likelihood and gradient calculation, and to perform optimization using LBFGS.

**Regularization**. For some cases the likelihood as a function of the overdispersion parameter can be extremely flat, leading to numerical instability. In order to stabilize the optimization we use very weak regularization in the form of the prior

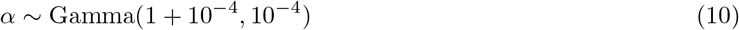

We experimented with two different versions of the *DM* GLM. The first uses a shared concentration parameter α*_j_* = α for all introns *j* in a cluster (the beta-binomial GLM is a special case of this model). The second allows a different α*_j_* for each intron in the cluster.

**Identifiability**. The *DM* GLM shares with the more standard Multinomial GLM that the form in Equation 2 has a spurious degree of freedom: in particular, adding a constant to the input of the softmax does not change its output. To remove this degree of freedom from the model we parameterize each *β_j_* as

**Figure S15:**
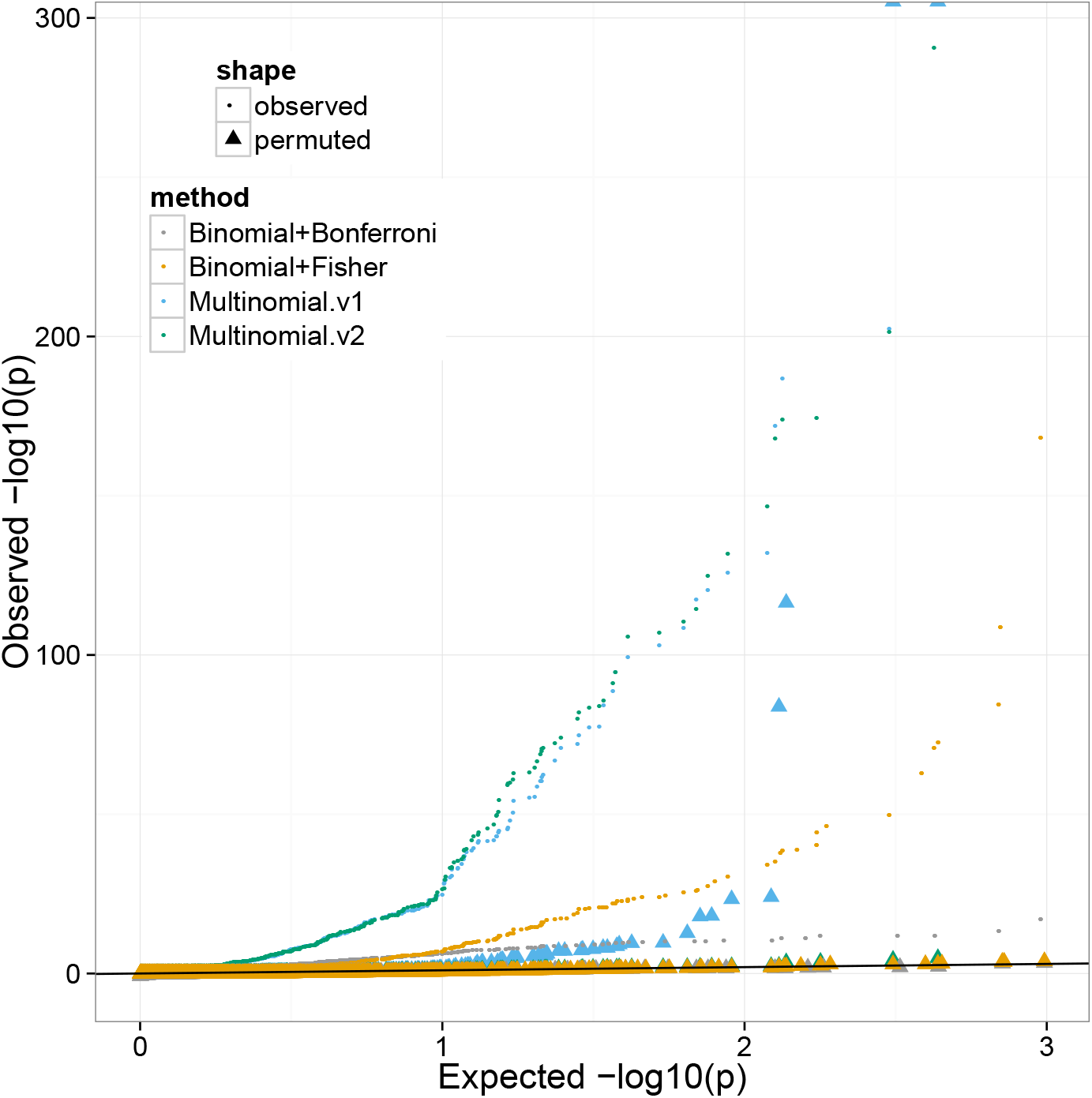
Comparison between beta-binomial and Dirichlet-multinomial models for differential splicing analyses, performed on 10 male brain vs. heart samples from GTEx. Two approaches for combining per-intron p-values into cluster level introns are compared: Bonferroni correction and Fisher’s combined test. Bonferroni is very conservative, as expected. Fisher’s combined test has considerably lower power than the multinomial approaches. However, only v2 of the Dirichlet-multinomial (which uses a per intron concentration/overdispersion parameter) is well calibrated under permutations.

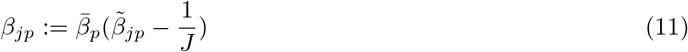

where 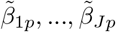 is constrained to lie on the *J*-simplex, i.e. 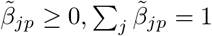, a constraint Stan naturally handles using a change of variables.

### 5.2 Likelihood ratio tests

Likelihood ratio tests are generally better calibrated than alternatives such as Wald statistics for testing for the significance of covariates, especially for modest sample sizes. We optimize wrt to *β, µ, α* separately for the null and alternative models (excluding and including the group indicator x respectively) to obtain log likelihoods *λ_0_* and *λ_1_* (for efficiency we initialize the optimization for the alternative model using the null model parameters) and then perform a likelihood ratio test: under the null 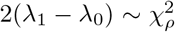 where *ρ* is the appropriate degrees of freedom. For the beta-binomial GLM *ρ* = *P_1_* – *P_0_* where *P_0_* and *P_1_* are the number of covariates in the null and alternative models respectively. For the Dirichlet-multinomial GLM we have *ρ* = (*J* – 1)(*P*_1_ – *P*_0_) where *J* is the number of introns in the cluster.

## 6 Differential intron excision analyses

### 6.1 Identification of tissue-dependent intron excision levels

We used LeafCutter’s Dirichlet-multinomial GLM to identify intron clusters with at least one differentially excised intron. We searched for intron excision level differences between all tissue pairs. However, we should note that owing to sample size differences, we will have different power to detect differential splicing of varying magnitude between pairs (we can detect splicing differences of small magnitude only in comparisons with large sample sizes). When we hierarchically clustered all samples according to the intron excision levels of introns that were present (i.e. were supported by reads) in all species, we saw a mix between tissue and species clustering (Figure S16). However, when we conditioned on introns that were differentially excised across human tissue pairs according to LeafCutter, we saw a clear clustering by tissue (Figure 2a).

**Figure S16:**
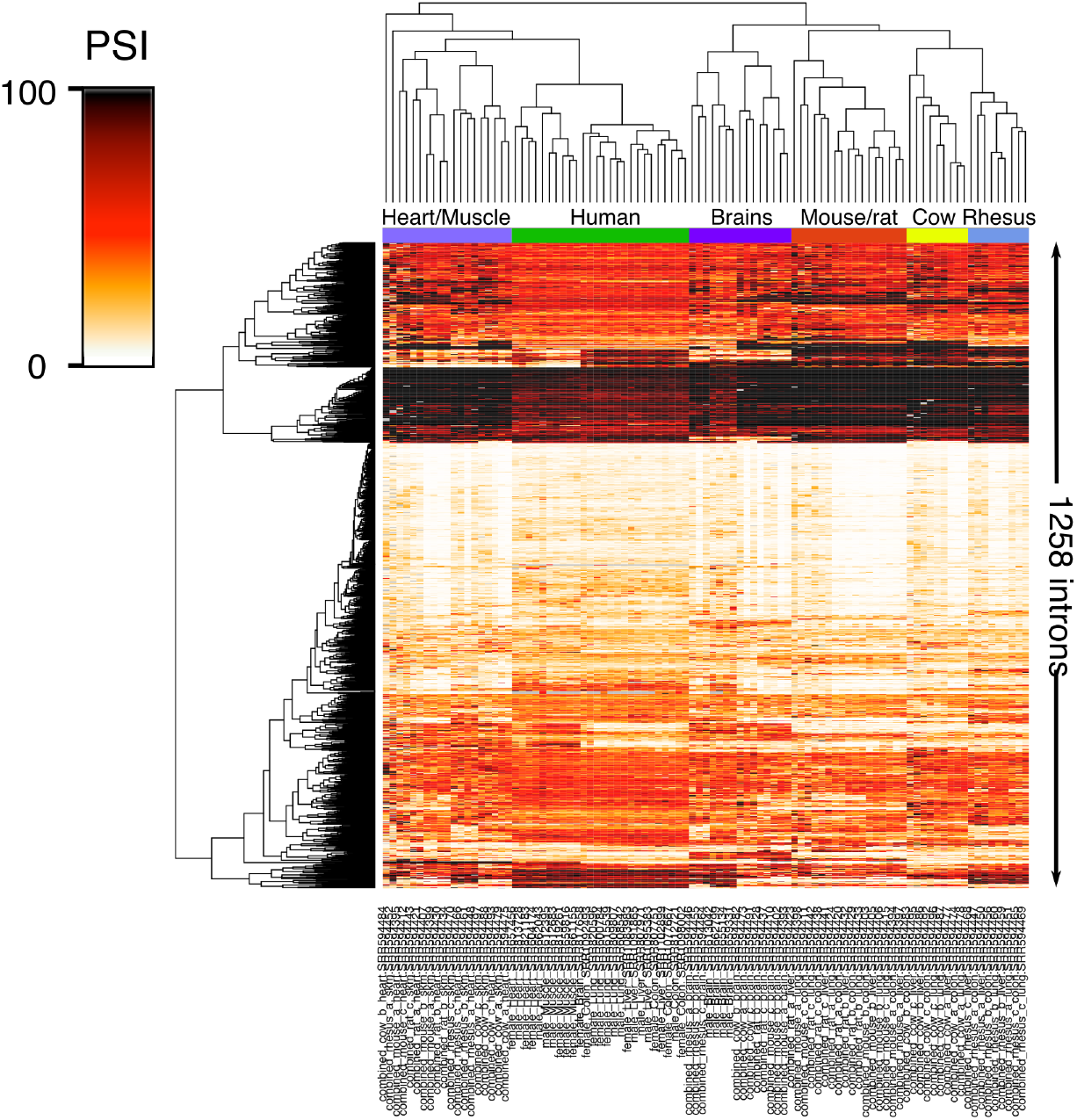
Hierarchical clustering on all 1,258 introns that had no missing values in any of the samples.

### 6.2 Effectiveness at small sample sizes

RNA-seq experiments are often performed on a handful of samples only. To determine whether LeafCutter is effective in this setting we performed clustering, quantification and differential intron usage analysis on 4 male brain and 4 male muscle samples from GTEx. As a “bronze standard” we additionally performed quantification and differential splicing on 110 muscle and 110 brain samples (using the introns and clusters identified using 8 samples). With only *N* = 8 samples, LeafCutter appears to be well-calibrated under permutations (Figure S17a) and has sufficient power to detect 885 clusters with evidence of differential intron usage (FDR 10%, maximum absolute effect size > 1.5), compared to 1906 found at *N* = 220. The per cluster *p*-values are highly correlated between the small and full sample sizes (R^2^ = 0.72, Figure S17b), and 98% of the clusters significant at *N* = 8 are also significant at *N* = 220. Per intron effect sizes between the two sample size settings are also highly correlated (R^2^ = 0.49, Figure S17c), although as expected the variance of the *N* = 8 effect sizes is large. This is particularly the case when the intron is only observed at all in one of the two tissues (Figure S17d).

### 6.3 Pan-mammalian tissue clustering of intron excision profiles

To evaluate the conservation of intron excision profiles across mammalian tissues, we used OLego to map RNA-seq data ^27^ from eight organs (testes, heart, kidney, liver, lung, brain, colon, and spleen) in four mammals (mouse, rat, cow, and rhesus macaque) to their respective genomes. We then projected all introns supported by RNA-seq reads onto the human genome using liftOver and clustered projected introns from all four mammals and human GTEx samples using LeafCutter. We then focused on four disjoint pairwise comparisons (Testis vs Kidney, Muscle vs Colon, Heart vs Lung, and Brain vs Liver, Figure S18).

**Figure S17:**
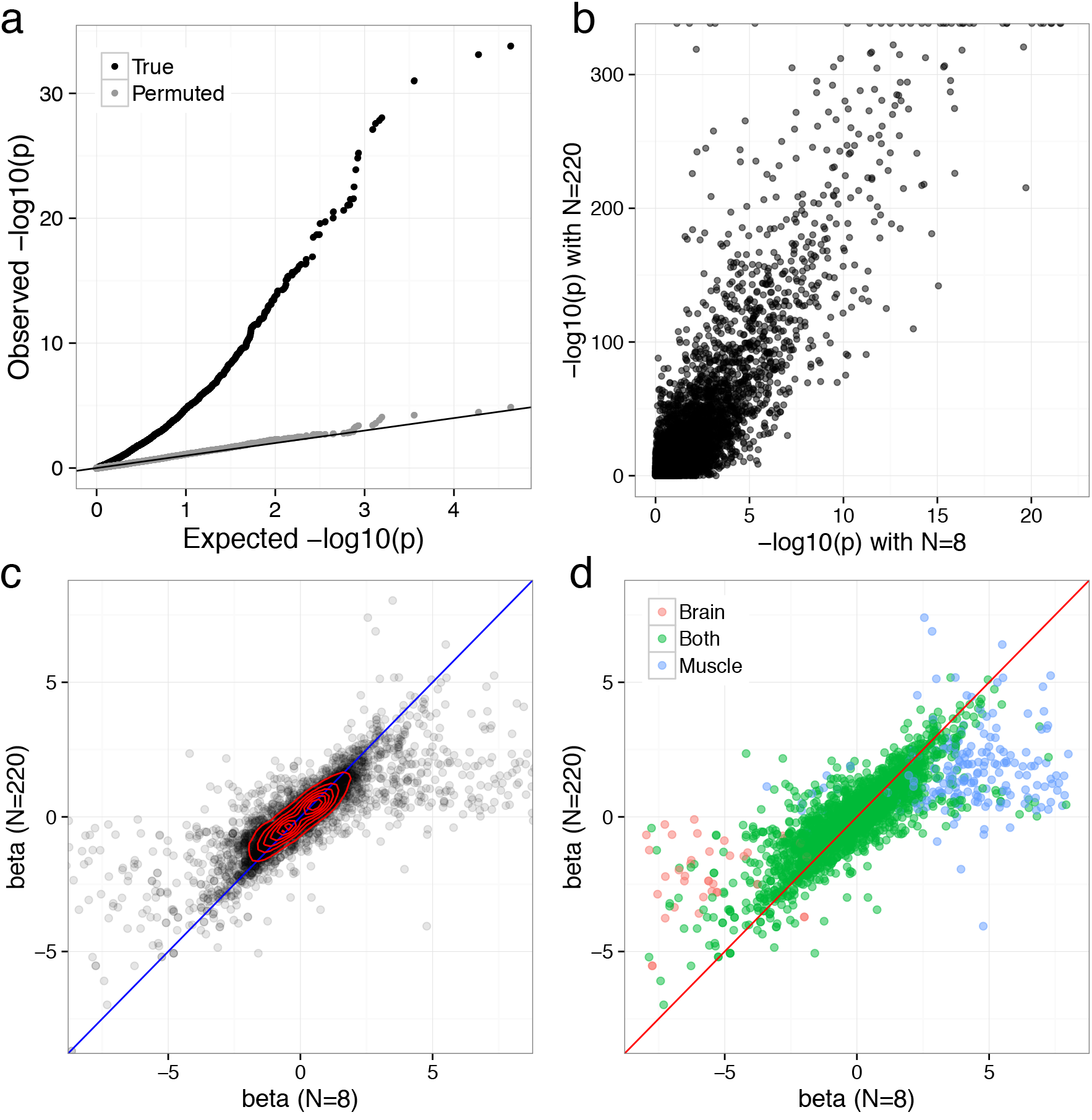
LeafCutter is effective even with as few as 8 samples. Here we performed differential splicing analysis of 4 male brain vs 4 male muscle samples, and compared to results using 220 samples. **a**) *p*-values under permutations are well-calibrated. **b-c**) *p*-values and effect sizes are highly correlated between the two sample size datasets. **d**) Significant disparity in effect sizes between the two sample sizes is primarily driven by an intron being unique to a tissue when *N* = 8.

**Figure S18:**
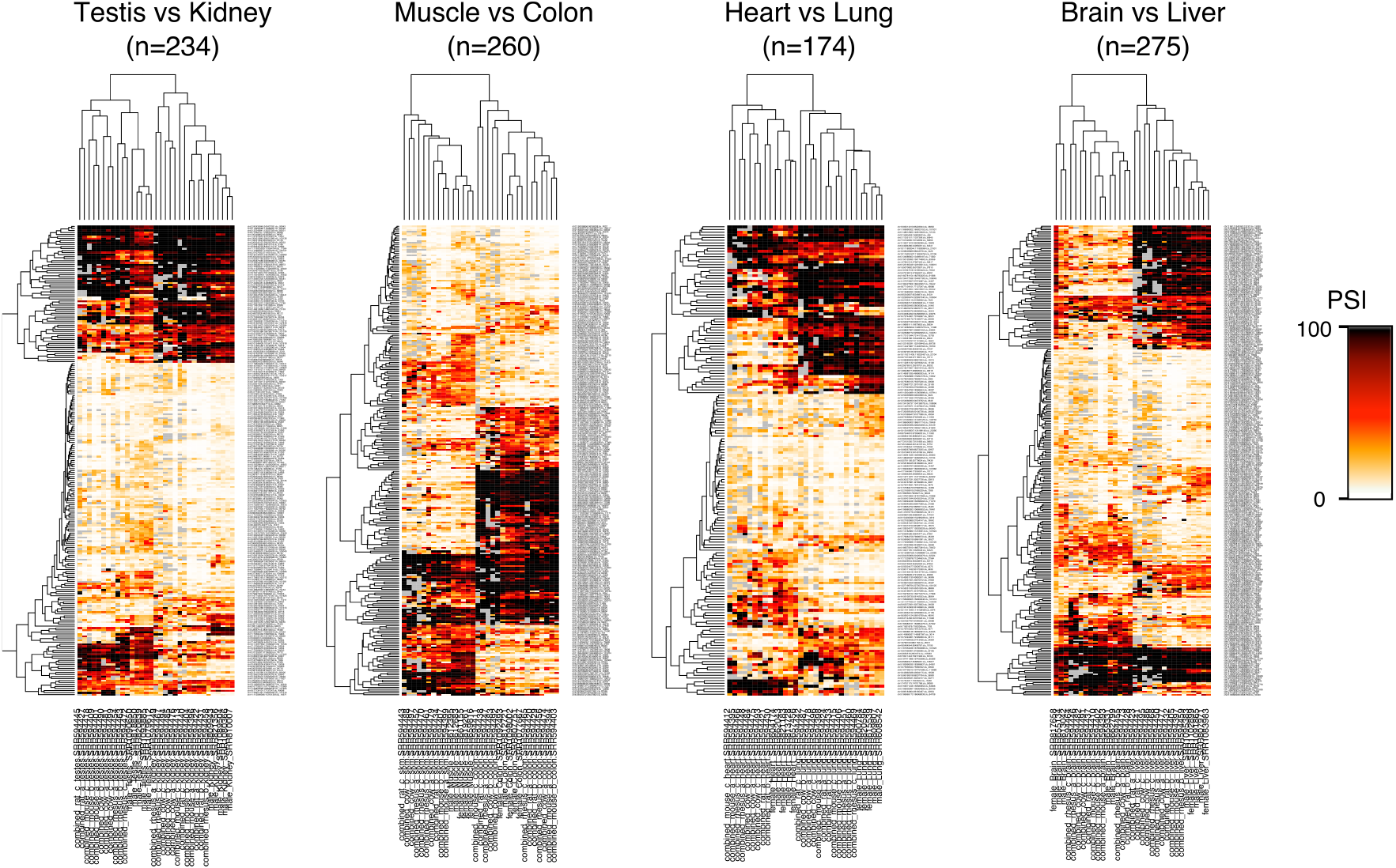
We restricted to introns that were found to be differentially excised between human tissues (p-value < 10^-10^ and effect size > 1.0)

## 7 sQTL mapping using LeafCutter

### 7.1 Mapping sQTLs in GEUVADIS LCL samples (linear regression)

To map sQTLs in GEUVADIS LCLs samples, we restricted our analysis on 372 samples derived from European individuals. We downloaded genotype files from ArrayExpress (E-GEUV-1). We used LeafCutter to obtain read proportions for all introns within alternatively excised intron clusters. We then standardized the values across individuals for each intron and quantile normalized across introns ^42^ and used this as our phenotype matrix. We then used linear regression (as implemented in fastqtl) ^30^ to test for associations between variants (MAF ≥ 0.05) within 100kb of intron clusters and the rows of our phenotype matrix that correspond to the introns within each cluster. As covariate, we used the first 3 principal components of the genotype matrix plus the first 15 principal components of the phenotype matrix. To estimate the number of sQTLs at any given false discovery rate (FDR), we used the correct p-values from fastqtl, and then used Bonferroni correction to control for the number of introns we test per cluster (note that this is conservative). We then use Benjamini-Hochberg to estimate the FDR (sample permutations show that our association *p*-values at this step are well calibrated).

Unlike for YRI RNA-seq data where we used WASP ^18^ to correct for biases in allelic reads, we did not correct for biases caused by allelic reads for the CEU comparisons to keep comparisons fair with previous GEUVADIS analyses. To avoid biases, we removed all associations that might be caused by SNPs that overlap junction reads. To do this, we removed all intron clusters that had a variant that were 70 or fewer base pairs (GEUVADIS RNA-seq read length is 75bp and at least 6nt must overlap with all exons) away from the splice sites (in the exonic part).

### 7.2 sQTL mapping comparison between LeafCutter, Cufflinks2 and Altrans

We ran LeafCutter, Cufflinks2 and Altrans to estimate isoform and splicing events usage, respectively, on all 85 Yoruba WASP-processed ^18^ RNA-seq aligned data. We then standardized the values across individuals for each isoform/splicing event usage and quantile normalized across introns ^42^. As covariate, we used the first 3 principal components of the genotype matrix plus the first 15 principal components of the phenotype matrix. We then used fastqtl ^30^ to test for associations between variants (MAF ≥ 0.05) within 50kb of the transcript (Cufflinks), the splicing event (Altrans), or splicing cluster (LeafCutter). To estimate the number of sQTLs at any given false discovery rate (FDR), we used the correct p-values from fastqtl, then use Benjamini-Hochberg to estimate the FDR. Altrans discovers splicing events using a forward and a reverse pass on the aligned RNA-seq data, thereby producing two measurement tables. To allow fair comparison between Altrans and the other methods, we combined forward and reverse splicing QTLs and collapsed all events that shared a splice site together (as is done in LeafCutter).

### 7.3 Sharing of sQTL discoveries between LeafCutter, Cufflinks2 and Altrans

To quantify the proportion of LCL sQTLs that are shared between Cufflinks2, Altrans, and LeafCutter, we first took the most significant SNP-gene/cluster pairs for every gene/clusters that had a sQTL at a 10% FDR. Note here that the following observations were qualitatively the same when we used a 1% FDR cutoff. We then collected the *p*-values of the associations of the SNP-gene pairs (when there were more than one splicing event tested per genes, we took the minimum *p*-values times the number of tested events) (Figure S19) and used the Storey’s π0 method ^43^ to estimate the proportion of shared discoveries (Figure S20).

**Figure S19:**
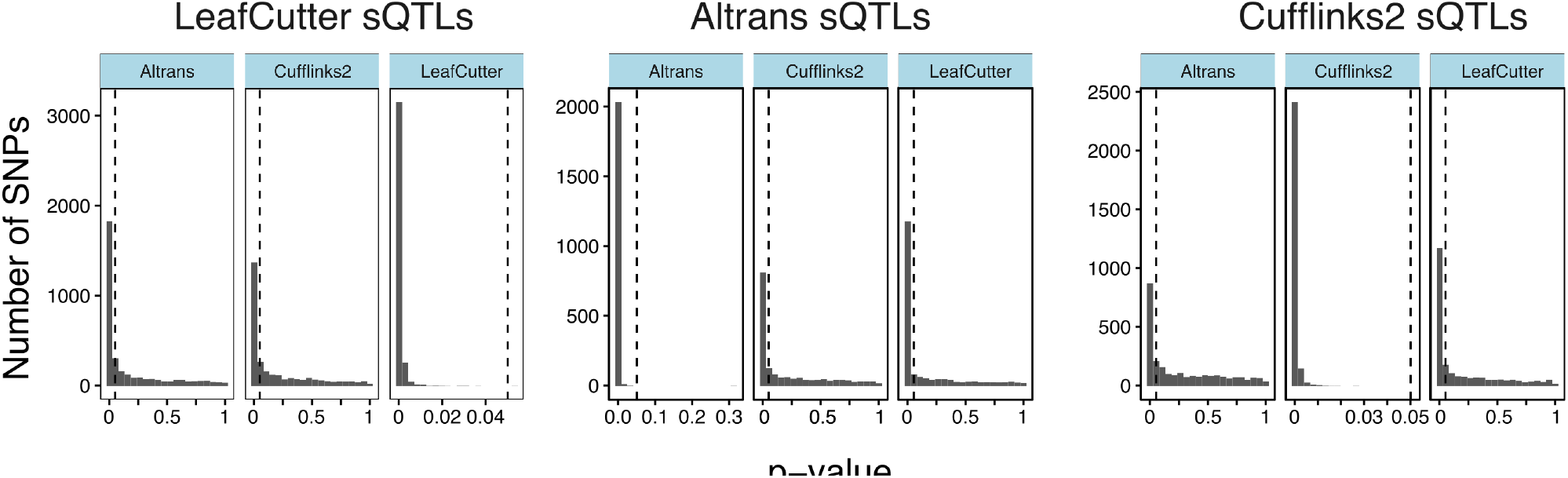
Distribution of SNP-gene splicing association p-values. Three panels correspond to sQTLs identified at 10% FDR using LeafCutter, Altrans, and Cufflinks2, respectively.

Overall, we find a higher pairwise sQTL sharing between LeafCutter and either of the two other methods (Altrans and Cufflinks2) than compared to the sharing between Altrans and Cufflinks2. Conversely, we found that while LeafCutter identified more sQTLs at 10% FDR, LeafCutter sQTLs were more enriched in low sQTL p-values as measured by Altrans or Cufflinks2. These observations suggest that LeafCutter is both more sensitive (lower proportion of false negatives) and more accurate (lower proportion of false positives).

### 7.4 Mapping sQTLs in GEUVADIS LCL samples (Dirichlet-multinomial GLM)

In addition to using linear regression, we also used LeafCutter’s Dirichlet-multinomial GLM to map sQTLs. This approach has two main advantages: (1) it accounts for the over-dispersion of read count data, and (2) it combines signal from changes in intron excision levels across the entire cluster instead of considering each intron independently. However, when we applied to our GEUVADIS data and controlled FDR using permutations, we found fewer sQTLs than our linear model approach, likely driven by clusters with heavy-tailed count distributions which are effectively handled by the quantile normalization in the linear approach.

**Figure S20:**
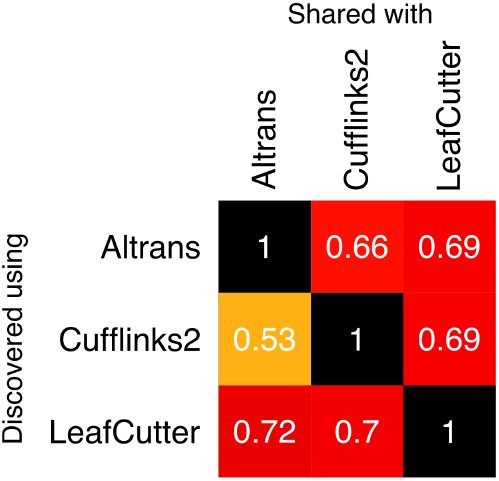
Sharing of sQTL discoveries between Cufflinks2, Altrans, and LeafCutter estimated using Storey’s π_0_ method.

### 7.5 Mapping sQTLs in four GTEx tissues

To identify sQTLs in GTEx tissues, we used the same strategy as in GEUVADIS LCLs (linear regression). However, we used the first 5 genotype PCs and the first 10 PCs as covariates (5+10 instead of 3+15).

**Table S20:** Sample sizes of processed GTEx .bam files for sQTL mapping.

| Tissue | Number of individuals |
| --- | --- |
| Heart | 95 |
| Blood | 170 |
| Lung | 128 |
| Thyroid | 118 |

## 8 sQTL analyses

### 8.1 Identification of functional enrichment of sQTLs

To identify functional categories enriched in sQTLs, we first annotated all variants using SnpEff version 4.1f. We next sampled at random ∼200,000 SNPs that are located near genes (i.e. had the annotation “Upstream”, “Downstream”, “Intronic”, or were exonic variants). This is because we only test SNPs that are near genes. The number of sampled SNPs corresponds to 50 times the number of sQTLs identified in our study. We computed the log-fold enrichment in functional annotations of the top most significant sQTLs (n = 4, 543) over this random sample of SNPs. Finally, to obtain confidence intervals, we repeated the random sampling procedure 500 times.

**Figure S21:**
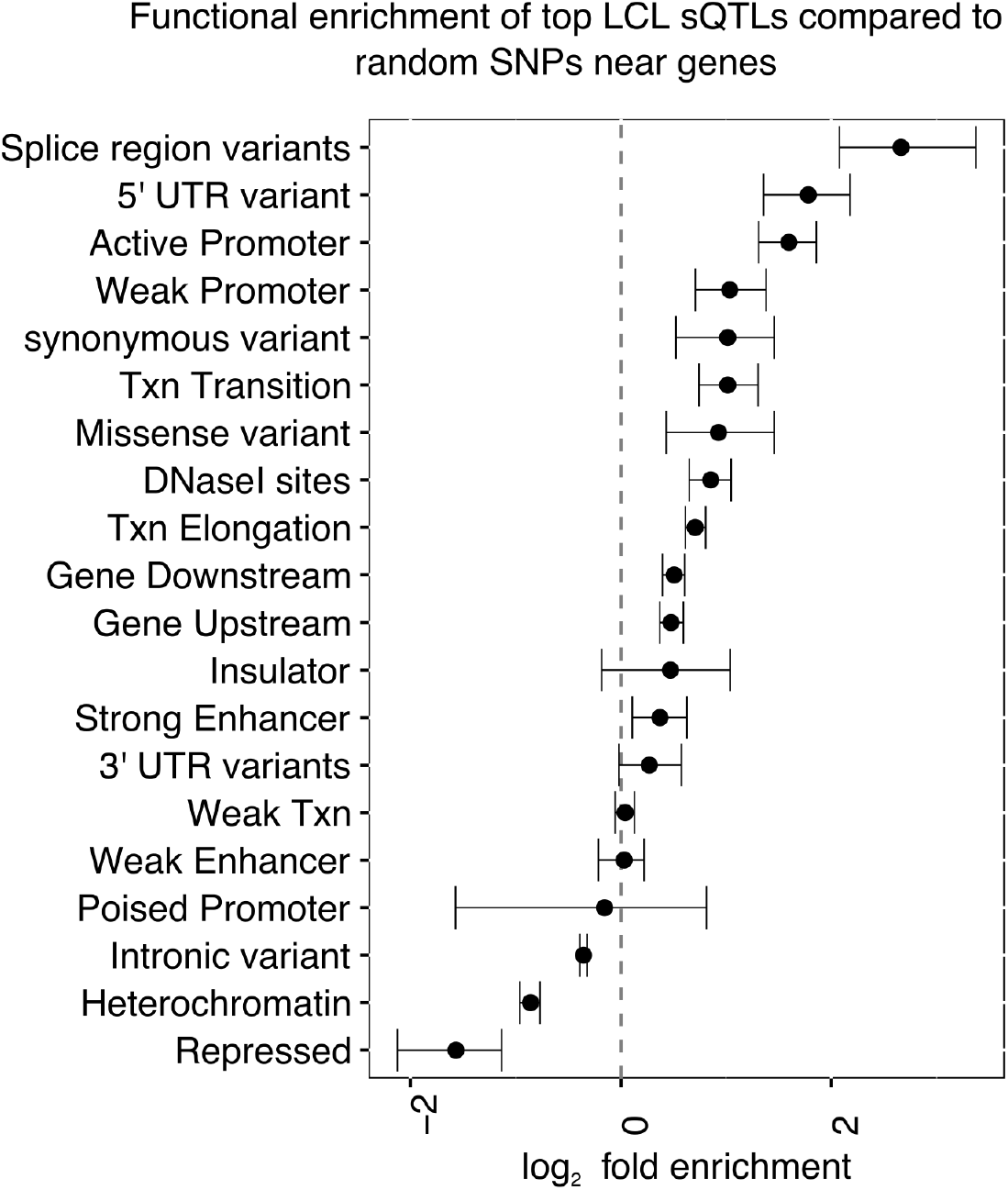
Functional enrichment of 4,543 sQTLs identified at 1%FDR from CEU GEUVADIS data. Bar represent 95% confidence interval from 500 bootstraps.

### 8.2 Comparison with GEUVADIS exon eQTLs, and trQTLs

Although LeafCutter does not explicitly search for genetic variants that are associated with differences in exon level splicing or transcript ratios, we expected that these variants will also affect intron excision, which are detected by LeafCutter. To verify this, we compared the distribution of p-values from the association between LeafCutter intron excision and genome-wide SNPs to the p-values from the association between LeafCutter intron excision and SNPs that were previously classified as exon eQTLs and transcription ratio QTLs in GEUVADIS. More specifically, we downloaded the list of exon eQTLs and trQTLs from ArrayExpress (E-GEUV-3) and for each exon/gene took the SNP with the strongest association to exon level or transcript ratio. We then computed the association p-values of these SNPs with all tested LeafCutter intron excision levels, using Bonferroni correction to adjust our p-values. As expected both exon eQTL and trQTL SNPs were enriched in strong associations to intron excision levels compared to random SNPs, and trQTL SNPs were most enriched in strong associations.

**Figure S22:**
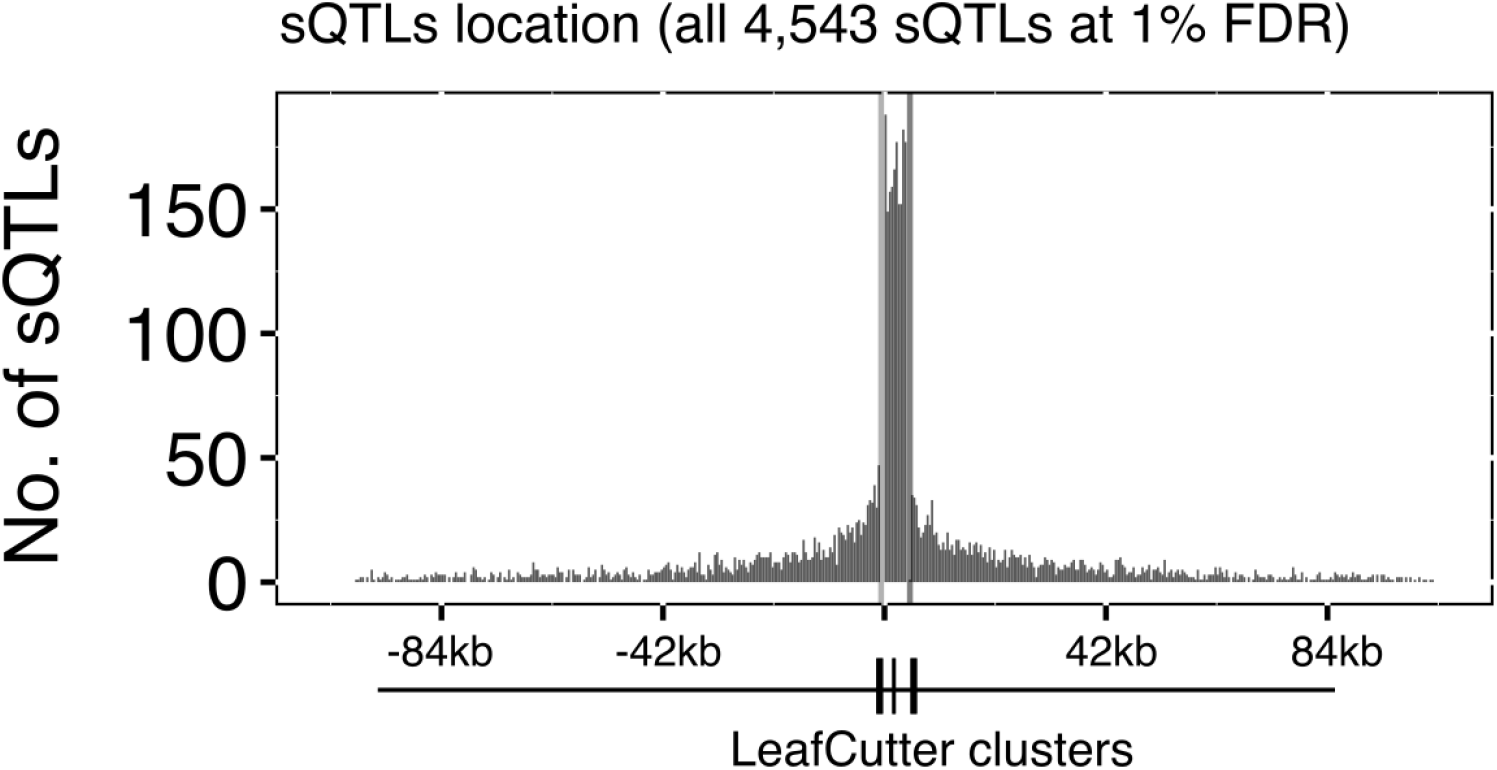
Meta-cluster representation of position of all 4,543 sQTLs identified at 1%FDR.

We next wished to verify that trQTLs detected in GEUVADIS were mostly identified as LeafCutter intron sQTLs. We again took the best trQTL SNP for each gene, and estimated the number that were associated with a cluster at a corrected *p*-value < 0.05. To correct for SNPs tested against multiple clusters, we used Bonferroni correction to adjust the *p*-value of the strongest association. We find that 399 (81.3%) of the 491 top trQTLs we tested are significantly associated (*p* < 0.05); this percentage is likely higher because our Bonferroni correction is conservative. Furthermore, as expected, when we use the same procedure to ask how many of the top 491 trQTLs are significantly associated to intron splicing when our sample labels are permuted, we find that only 4.7% are (our statistical tests are well calibrated; ∼5% of our tests should achieve a 0.05 significance under the null model).

### 8.3 Relationship between gene expression levels and power to detect sQTLs

We examined the expression profiles of the genes with significant sQTLs detected by LeafCutter. As expected we found a strong positive relationship between our power to detect a sQTL for a gene and the expression level of a gene (Figure S23a). Indeed, while most annotated genes (including non protein-coding genes) were expressed at very low levels, we found almost no sQTLs for genes whose expression were less than 0.025 RPKM. While there is a clear decrease in LeafCutter’s ability to identify sQTLs in lowly expressed genes (Figure S23a), we were able to find sQTLs for many lowly-expressed genes, starting from 0.1 RPKM (Figure S23b).

**Figure S23:**
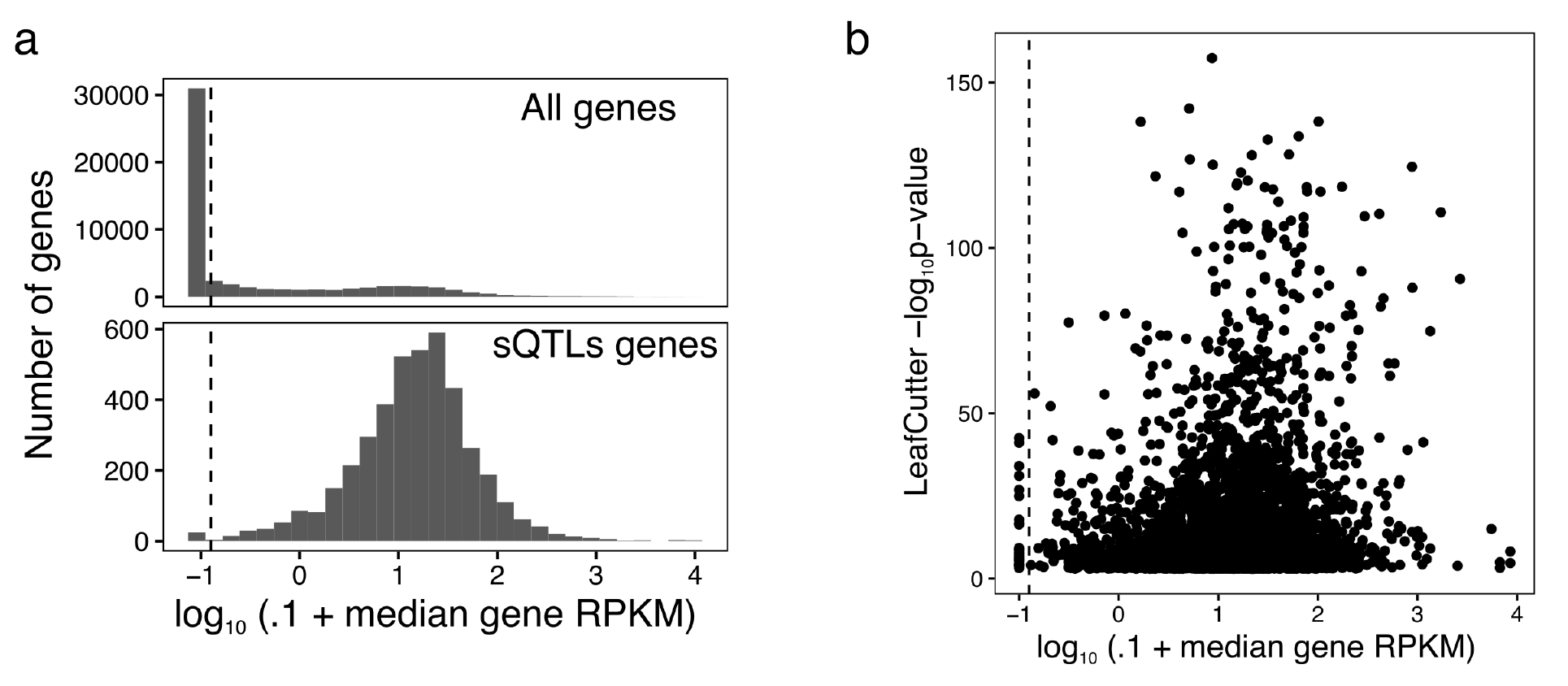
(a) Distribution of median LCLs gene expression levels for all genes (top) and genes with one or more LeafCutter sQTLs. (b) Scatter plot of LeafCutter p-value associations with respect to the expression levels of the corresponding genes. Dashed lines correspond to approximately 0.025 RPKM.

### 8.4 Replication of sQTLs across GTEx tissues

To estimate the proportion of sQTLs that are replicable across tissue types, we took the best SNP of each sQTL-cluster pair for each tissue and asked whether the sQTL association was significant (*p* < 0.05) in another tissue. This estimate is likely to be conservative as it does not account for incomplete power. The replication is therefore likely to be even higher than our current estimates of 75–93%.

**Figure S24:**
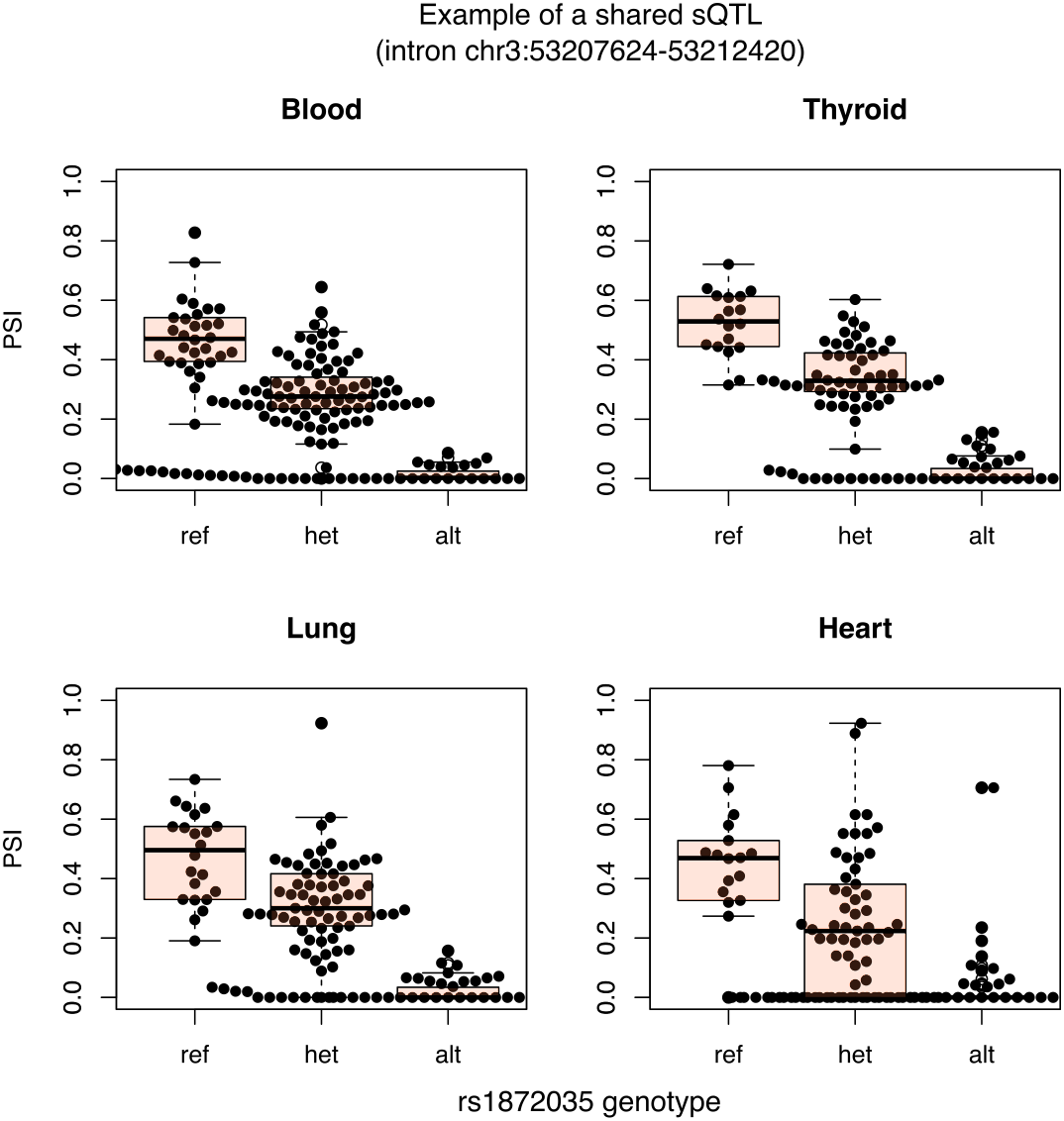
Example of a shared sQTL.

### 8.5 Tissue-specific sQTLs

To identify tissue-specific sQTLs, we searched for genetic variants that were associated significantly with intron excision levels in one tissue, but not in any of the other three tissues (*p* > 0:1), requiring all tissues to have junction reads in the intron cluster.

**Figure S25:**
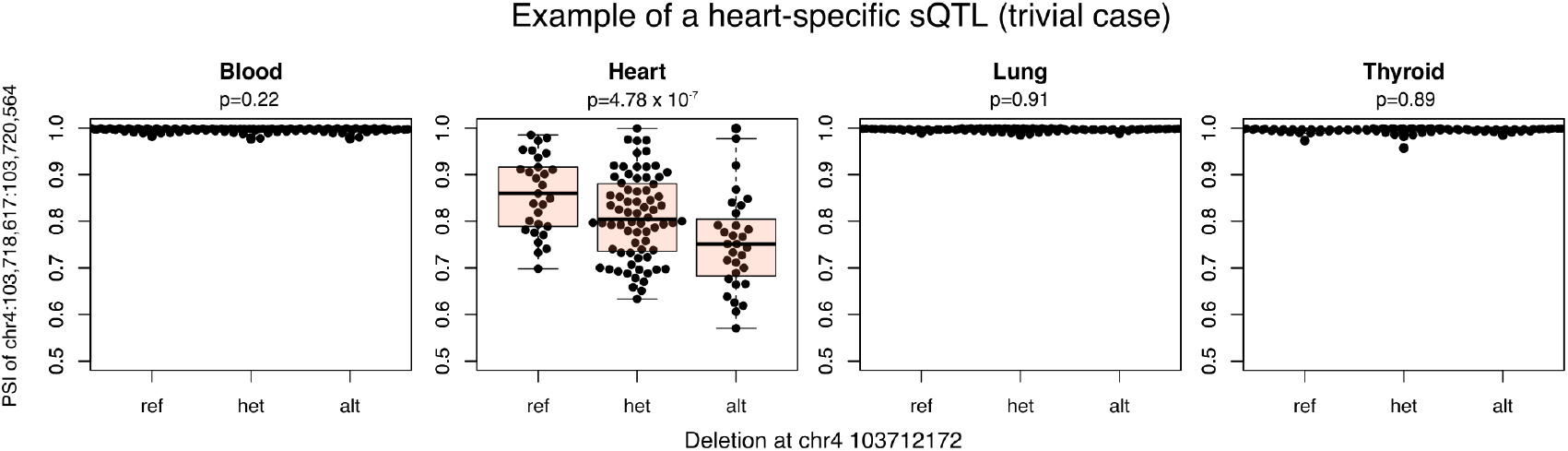
Example of a tissue-specific sQTL.

## 9 LeafCutter sQTL signals in genome-wide association studies

To verify that LeafCutter sQTLs can help us identify disease-associated variants that function by modulating splicing, we downloaded summary statistics from two autoimmune GWAS studies (multiple sclerosis ^44^ and rheumatoid arthritis ^45^) and looked for enrichment of strong association *p*-values among the top LeafCutter sQTLs and GEUVADIS gene eQTLs (we removed the extended MHC region from this analysis). We found that 1,205 LeafCutter sQTL SNPs and 901 GEUVADIS eQTL SNPs (the SNP with most significant p-value) were also tested (with >5% MAF) in the multiple sclerosis genome-wide association study, and that 3,069 LeafCutter sQTL SNPs and 2,250 GEUVADIS eQTL SNPs were tested in the rheumatoid arthritis study. We then took the QTLs and plotted the distribution of – log_10_ (p-value) of their association to each trait separately. As expected ^17^, we found that LeafCutter sQTLs were more highly enriched in associations with low p-values compared to GEUVADIS eQTLs in multiple sclerosis and were similarly enriched in rheumatoid arthritis. This is notable because we considered a larger number of LeafCutter sQTLs than GEUVADIS eQTLs for both diseases. These observations suggest that LeafCutter allows us to identify as many or more disease-associated variants that *act* by affecting splicing as compared to those that *act* by affecting total expression levels.

### 9.1 Prediction Models and S-PrediXcan

Prediction models were trained by fitting Elastic-Net linear models to each gene for the expression models and to each intron cluster for the splicing models using nearby SNPs dosages as features. Before fitting the models, we removed non biallelic SNPs and any ambiguously stranded SNPs from the genotype data. We downloaded normalized and PEER corrected expression data from the GEUVADIS study. Intron excision traits were corrected for genetic principal components and covariates (as outlined above). Once the data had been preprocessed, for each gene or intron cluster, SNPs within 1Mb upstream and 1Mb downstream of their start and end sites were selected as variables for the model. Using the R package glmnet we fit a 10-fold cross-validated Elastic-Net linear model using a mixing parameter of 0.5 for each gene and intron cluster. Further details can be found in ^36,33,37^ and training pipelines can be downloaded from github.com/hakyimlab/PredictDBPipeline.

A total of 4625 gene associations were obtained for the genetic expression model, and 41196 intron quantification cluster associations for the splicing model, that had a model prediction FDR < 5% (computed from the correlation between cross validated prediction and observed values).

**Table S25:**
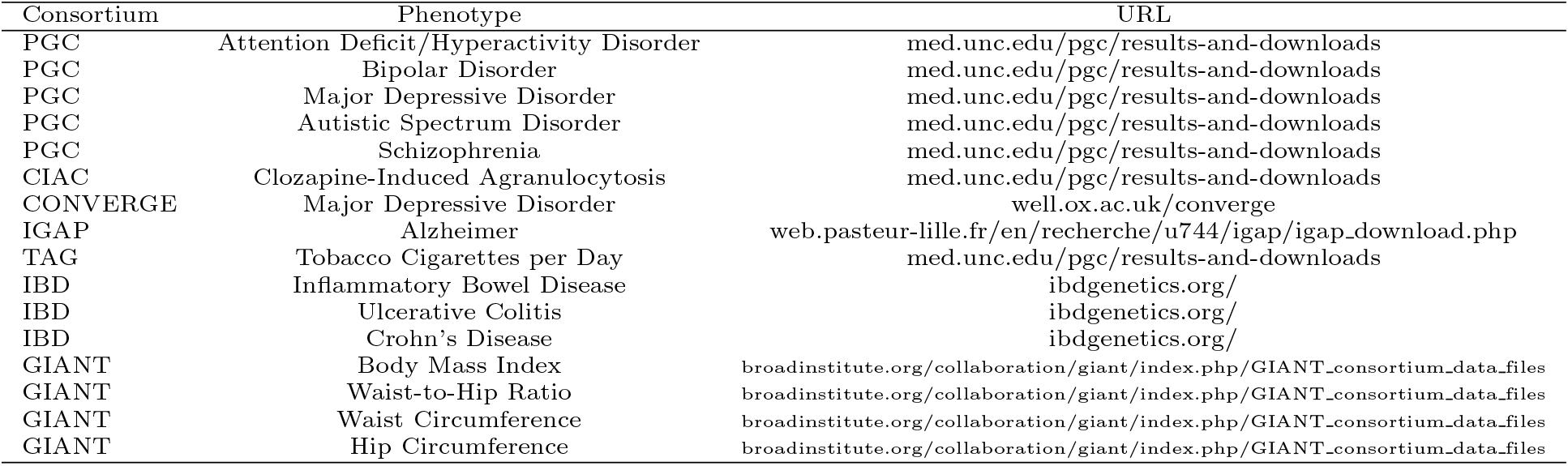
List of Genome-wide Association Meta Analysis (GWAMA) Consortia and phenotypes.

| Consortium | Phenotype | URL |
| --- | --- | --- |
| PGC | Attention Deficit/Hyperactivity Disorder | med.unc.edu/pgc/results-and-downloads |
| PGC | Bipolar Disorder | med.unc.edu/pgc/results-and-downloads |
| PGC | Major Depressive Disorder | med.unc.edu/pgc/results-and-downloads |
| PGC | Autistic Spectrum Disorder | med.unc.edu/pgc/results-and-downloads |
| PGC | Schizophrenia | med.unc.edu/pgc/results-and-downloads |
| CIAC | Clozapine-Induced Agranulocytosis | med.unc.edu/pgc/results-and-downloads |
| CONVERGE | Major Depressive Disorder | well.ox.ac.uk/converge |
| IGAP | Alzheimer | web.pasteur-lille.fr/en/recherche/u744/igap/igap_download.php |
| TAG | Tobacco Cigarettes per Day | med.unc.edu/pgc/results-and-downloads |
| IBD | Inflammatory Bowel Disease | ibdgenetics.org/ |
| IBD | Ulcerative Colitis | ibdgenetics.org/ |
| IBD | Crohn's Disease | ibdgenetics.org/ |
| GIANT | Body Mass Index | broadinstitute.org/collaboration/giant/index.php/GIANT_consortium_data_files |
| GIANT | Waist-to-Hip Ratio | broadinstitute.org/collaboration/giant/index.php/GIANT_consortium_data_files |
| GIANT | Waist Circumference | broadinstitute.org/collaboration/giant/index.php/GIANT_consortium_data_files |
| GIANT | Hip Circumference | broadinstitute.org/collaboration/giant/index.php/GIANT_consortium_data_files |

We downloaded genomewide association meta analysis (GWAMA) results for 40 phenotypes from 18 consortia and performed S-PrediXcan analysis using both expression and intron models. The full list of traits and consortia is displayed in Supplementary Table S25.

## 10 Processed data availability

**Figure S26:**
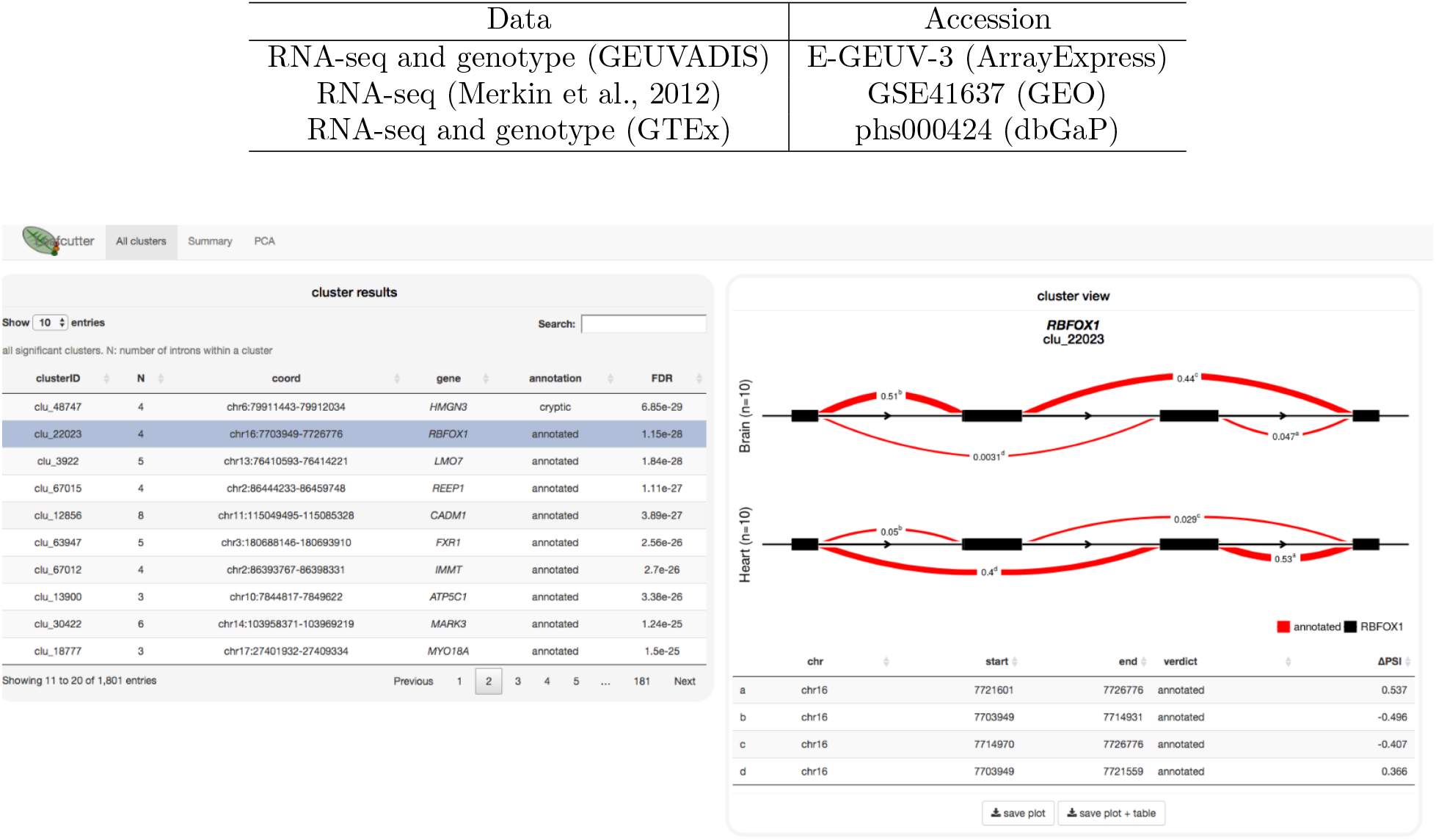
Visualization of differential splicing between heart and brain identified by comparing 10 GTEx heart and brain samples using LeafCutter. All cluster figures are available at https://leafcutter.shinyapps.io/leafvis2/.

